# The filipodia-like protrusions of adjacent somatic cells shape the developmental potential of mouse oocytes

**DOI:** 10.1101/2022.09.15.508092

**Authors:** Flora Crozet, Gaëlle Letort, Christelle Da Silva, Adrien Eichmuller, Anna Francesca Tortorelli, Morgane Belle, Julien Dumont, Tristan Piolot, Aurélien Dauphin, Fanny Coulpier, Alain Chédotal, Jean-Léon Maître, Marie-Hélène Verlhac, Hugh.J Clarke, Marie-Emilie Terret

## Abstract

The oocyte must grow and mature before fertilization, thanks to a close dialogue with the somatic cells which surround it. Part of this communication is through filopodia-like protrusions, called transzonal projections (TZPs), sent by the somatic cells to the oocyte membrane. To investigate the contribution of TZPs to oocyte quality, we impaired their structure by generating a full knockout mouse of the TZP structural component Myosin-X (MYO10). Using spinning disk and super-resolution microscopy combined with a machine learning approach to phenotype oocyte morphology, we show that the lack of *Myo10* decreases TZP density during oocyte growth. Reduction in TZPs does not prevent oocyte growth but impairs oocyte-matrix integrity. Importantly, we reveal by transcriptomic analysis that gene expression is altered in TZP-deprived oocytes, and that oocyte maturation and subsequent early embryonic development are partially affected, effectively reducing mouse fertility. We propose that TZPs play a role in the structural integrity of the germline-somatic complex, which is essential for regulating gene expression in the oocyte and thus its developmental potential.

## Introduction

During oogenesis, mammalian oocytes undergo a process of differentiation that determines their quality and thus their developmental potential as early embryos after fertilization. Oocyte differentiation begins with the entry into a growth phase during which the oocyte increases in size and stores large amounts of macromolecules necessary to complete its development and early embryogenesis. During growth, the oocyte remains arrested in prophase and gradually acquires the competence to resume meiosis^1^. Once fully grown, it further differentiates by undergoing meiotic maturation, a process consisting of two successive meiotic divisions (meiosis I and II) generating a mature oocyte ready to be fertilized.

The oocyte does not progressively acquire meiotic and developmental competence by itself in an autonomous way, but through a close dialogue with the somatic cells that surround it, called follicular cells, which themselves undergo a parallel and intricate process of differentiation^2–5^. Together with the oocyte, the follicular cells form a germ-somatic complex called the ovarian follicle.

While the oocyte communicates with its surroundings via secreted factors^2,4,6^, somatic communication occurs mostly through direct cell-cell contacts mediated by actin-rich filopodia-like^7^ protrusions called transzonal projections (TZPs)^6,8–10^. These TZPs cross the *zona pellucida*, the extracellular matrix surrounding the oocyte, and contact the oocyte membrane at their tips. They allow follicular cells to metabolically support the oocyte and prevent early meiotic resumption^3,6^. In addition, TZPs may provide structural support in the follicle by maintaining cohesion between the oocyte and the follicular cells^9,11^.

While dialogue within the ovarian follicle is increasingly emerging as a major player in the emergence of oocyte quality, the role of direct cell-cell contact communication between the oocyte and the follicular cells is still poorly understood, and may shed additional light on the acquisition of oocyte developmental competence. In this study, we sought to characterize the contribution of TZPs to oocyte development by impairing the structure of TZPs. As a candidate gene, we selected Myosin-X (MYO10), a structural component of TZPs^12^ known to promote filopodia formation^13–15^ and potentially involved in the formation or maintenance of TZPs^16^. We generated a full knockout mouse of *Myo10* in all cell types to completely deplete it from the ovarian follicle (*Myo10*^-/-^ full). Furthermore, we took advantage of our previously generated mouse strain conditionally deleted for *Myo10* specifically in oocytes from early growth on (*Myo10^-/-^* oo)^17^ to help distinguishing phenotypes caused by loss of somatic MYO10 or loss of oocyte MYO10. We show that the loss of somatic but not oocyte MYO10 greatly decreases the density of TZPs. Surprisingly, oocytes deprived of TZPs develop to a normal size, but display oocyte-matrix defects. Importantly, gene expression is altered in TZP-deprived oocytes, correlating with lower rates of oocyte maturation and subsequent early embryonic development, inducing lower fertility in mice.

## Results

### Global deletion of *Myo10* decreases the density of transzonal projections without altering ovarian follicular organization

To deplete TZPs, we generated full knockout mice lacking the TZP component MYO10 in all cell types (Fig.S1 A-D). Loss of MYO10 expression in mice was previously characterized as semi-lethal, some animals dying during embryogenesis while others being able to develop until adulthood^15^. Similar to this study, we obtained adult *Myo10*^-/-^ mice (named *Myo10*^-/-^ full thereafter) and most showed phenotypes characteristic of the earlier described *Myo10* full knockout, such as the presence of a white spot on the abdomen and webbed fingers^15^ (Fig.S1 B). In addition to this strain, we used our previously generated mouse strain conditionally deleted for *Myo10* in oocytes from early growth on (named *Myo10*^-/-^ oo thereafter) as a control to potentially distinguish oocyte phenotypes related to germline loss of MYO10 versus those of somatic loss of MYO10^17^ (Fig.S1 A, D).

In control follicles, MYO10 is found diffuse in the cytoplasm of the oocyte and of follicular cells^17^ (Fig.1 A, S1 D) and accumulates into foci within the *zona pellucida* where TZPs are located^12,16^ (Fig.1 A, B). These foci were entirely lost from the *zona pellucida* of oocytes coming from *Myo10*^-/-^ full follicles (Fig.1 A, B, upper panels). Conversely, the foci were present in oocytes coming from *Myo10*^-/-^ oo follicles lacking MYO10 only in the oocyte (Fig.1 A, B, lower panels), indicating that MYO10 foci in the *zona pellucida* originate from follicular cells. Importantly, the loss of MYO10 foci in the *zona pellucida* of oocytes coming from *Myo10*^-/-^ full follicles was concomitant with a significant decrease in the density of TZPs (Fig.1 A, B, upper panels). This decrease was most apparent when imaging TZPs by super-resolution microscopy in *Myo10*^-/-^ full oocytes cleared of follicular cells (Movie.S1 OMX; Fig.1 C STED). This reduction was not observed in oocytes coming from *Myo10*^-/-^ oo follicles (Fig.1 A, B, lower panels), implying that TZP density is related to the presence of MYO10 in follicular cells.

**Figure 1:**
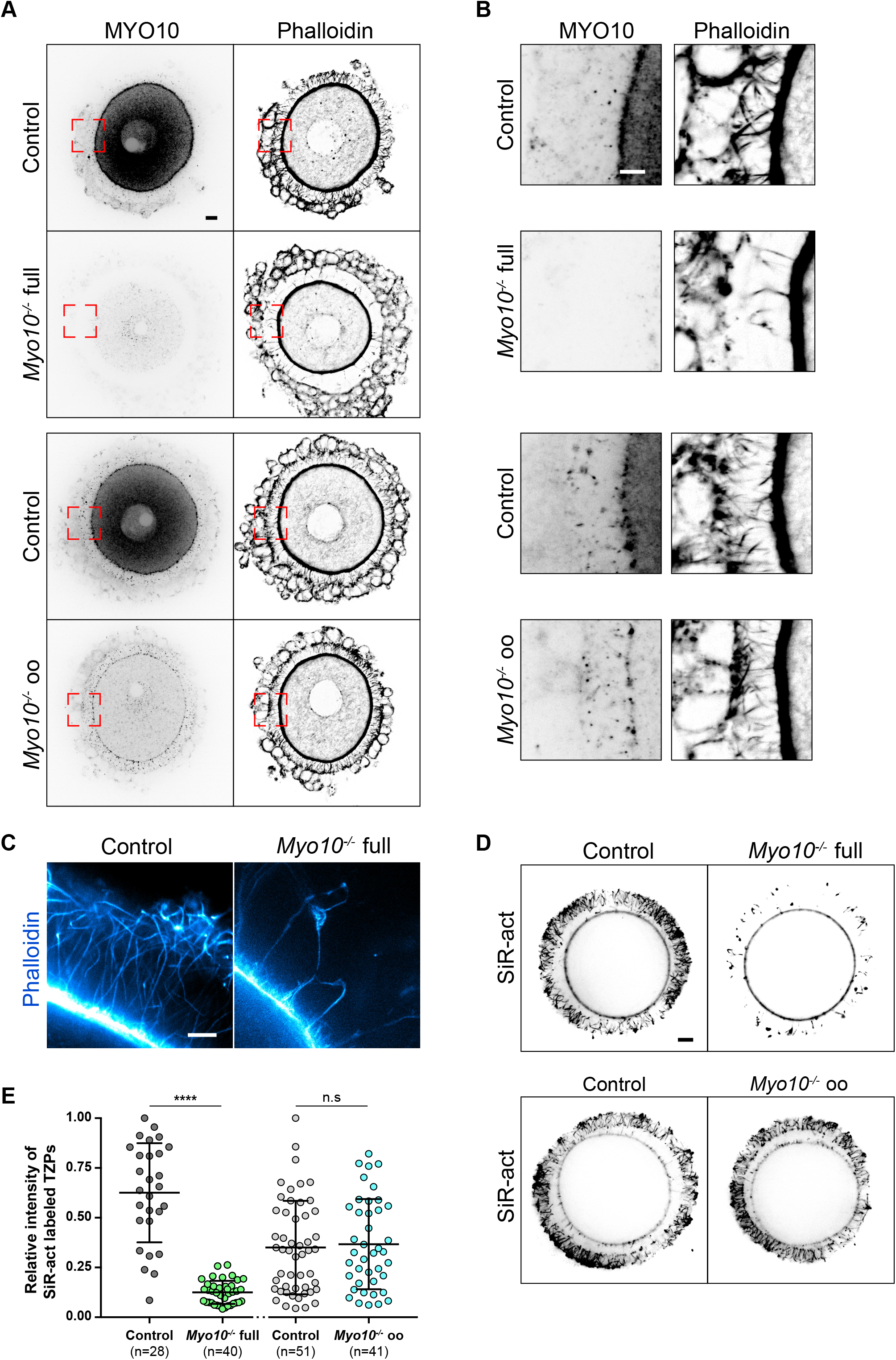
Global deletion of *Myo10* decreases the density of transzonal projections. **A)** Cumulus-oocyte complexes stained for Myosin-X (MYO10, left images) and with phalloidin to label F-actin (right images). The upper panel shows complexes from the *Myo10* full knockout strain (full) and the lower panel complexes from the *Myo10* oocyte knockout strain (oo). For each panel, control complexes are at the top and *Myo10*^-/-^ complexes at the bottom. For MYO1O staining, contrast adjustment is similar between complexes of the same strain. Scale bar 10 μm. **B)** Cropped images of the red dotted rectangles shown in A) focusing on the *zona pellucida*. MYO10 staining is on the left and phalloidin on the right. Scale bar 5 μm. **C)** STED microscopy images of phalloidin labeled oocytes arrested in prophase freed of follicular cells, focusing on the *zona pellucida* of a control oocyte (left) and a *Myo10*^-/-^ full oocyte (right). Scale bar 2 μm. **D)** Live fully grown oocytes arrested in prophase stained with SiR-actin (SiR-act) to label F-actin. The upper images are oocytes from the *Myo10* full knockout strain (full) and the lower images from *Myo10* oocyte knockout strain (oo). Controls are on the left and *Myo10*^-/-^ oocytes on the right. Scale bar 10 μm. **E)** Scatter plot of the intensity of all SiR-act labeled TZPs of fully grown oocytes arrested in prophase. Control and *Myo10*^-/-^ full oocytes are in dark gray and green, respectively. Control and *Myo10*^-/-^ oo oocytes are in light gray and blue, respectively. (n) is the number of oocytes analyzed. Data are mean±s.d. with individual data points plotted. Data are from three to five independent experiments. Statistical significance of differences was assessed by a two-tailed unpaired t test with Welch’s correction, P value < 0.0001 (*Myo10*^-/-^ full oocytes compared to control oocytes) and by a two-tailed unpaired t test, P value = 0.7271 (*Myo10*^-/-^ oo oocytes compared to control oocytes). n.s not significant.

Consistent with these observations in fixed follicles, we also measured a fivefold decrease in TZP density in live, fully grown *Myo10*^-/-^ full oocytes stained with SiR-actin to label F-actin (Fig.1 D, quantification in E). Conversely, TZP density remained similar to controls in live *Myo10*^-/-^ oo oocytes labeled with SiR-actin (Fig.1 D, E), indicating that *Myo10*^-/-^ oo oocytes retain a canonical density of follicular cell contacts. Since a few TZPs in the ovarian follicle do not contain actin^10,12^, we confirmed that loss of MYO10 not only decreases actin-rich TZPs but also total TZP density by staining oocytes with the membrane probe FM 1-43 (Fig.S2 A). We then wanted to determine when this phenotype of decreased TZP density appeared during follicular growth. Follicles at an early stage of growth contain few TZPs^12^, so it was difficult to determine whether there is a decrease in TZP density in *Myo10*^-/-^ full oocytes at the beginning of growth compared to controls. However, the phenotype of decreased TZP density in absence of MYO10 was apparent as early as mid-growth^16^ (Fig.S1 D right panel, magnifications), and TZP density was low in oocytes from *Myo10*^-/-^ full follicles at the end of growth regardless of the oocyte diameter (Fig.S2 B).

Impaired intercellular communication between the oocyte and follicular cells in mice deficient for the gap junction subunit connexin 37^18^ or deficient for the oocyte-secreted factor GDF9-20 led to ovaries with abnormal morphology, lacking fully grown follicles. We therefore tested whether decreasing TZP density altered the integrity of *Myo10*^-/-^ full ovaries. To gain access to intra-ovarian organization, whole ovaries from adult mice were transparized^21^. As in control ovaries, *Myo10*^-/-^ full ovaries contained late antral follicles (Fig.S3 A). *Myo10*^-/-^ full mice also showed a variation in the volume of their ovaries, (Movie.S2 A, B). Because of this variation, we assessed global ovarian integrity by measuring the follicular density of the ovary (number of follicles normalized by the ovarian area) rather than the absolute number of follicles per ovary. Follicular density was not significantly different between control and *Myo10*^-/-^ full ovaries (Fig.S3 B), indicating that the complete loss of MYO10 and subsequent reduction in TZP density do not alter global follicular organization in the ovary. Thus, we confirmed that MYO10 is a structural component of TZPs, essential for TZP formation or maintenance^12,16^.

### Oocytes deprived of TZPs reach a canonical size at the end of growth but are morphologically different from controls

To determine the TZP-mediated contribution of follicular cells to oocyte development, we assessed whether *Myo10*^-/-^ full oocytes deprived of TZPs developed properly. We used our previously developed deep-learning approach to characterize in an automatic manner the morphology of *Myo10*^-/-^ full oocytes at the end of the growth phase with the software Oocytor^22^ to assess their meiotic competence (defined as the ability of oocytes to resume meiosis at the end of growth and mature properly). First, *Myo10*^-/-^ full oocytes grew to normal sizes (Fig.2 A, B), and were arrested in prophase. We then measured the thickness and the perimeter of the *zona pellucida,* features that reflect the ability of the oocyte to resume meiosis^22^, and showed that they are similar between *Myo10*^-/-^ full oocytes and controls (Fig.S4 A-B). However, the texture of the *zona pellucida,* one of the two most important features to assess oocyte maturation potential^22^, was lower in *Myo10*^-/-^ full oocytes (Fig.2 C). Thus, the morphology of *Myo10*^-/-^ full oocytes at the end of growth suggests that they have a similar potential for meiosis resumption as controls, but a decreased potential for oocyte maturation.

**Figure 2:**
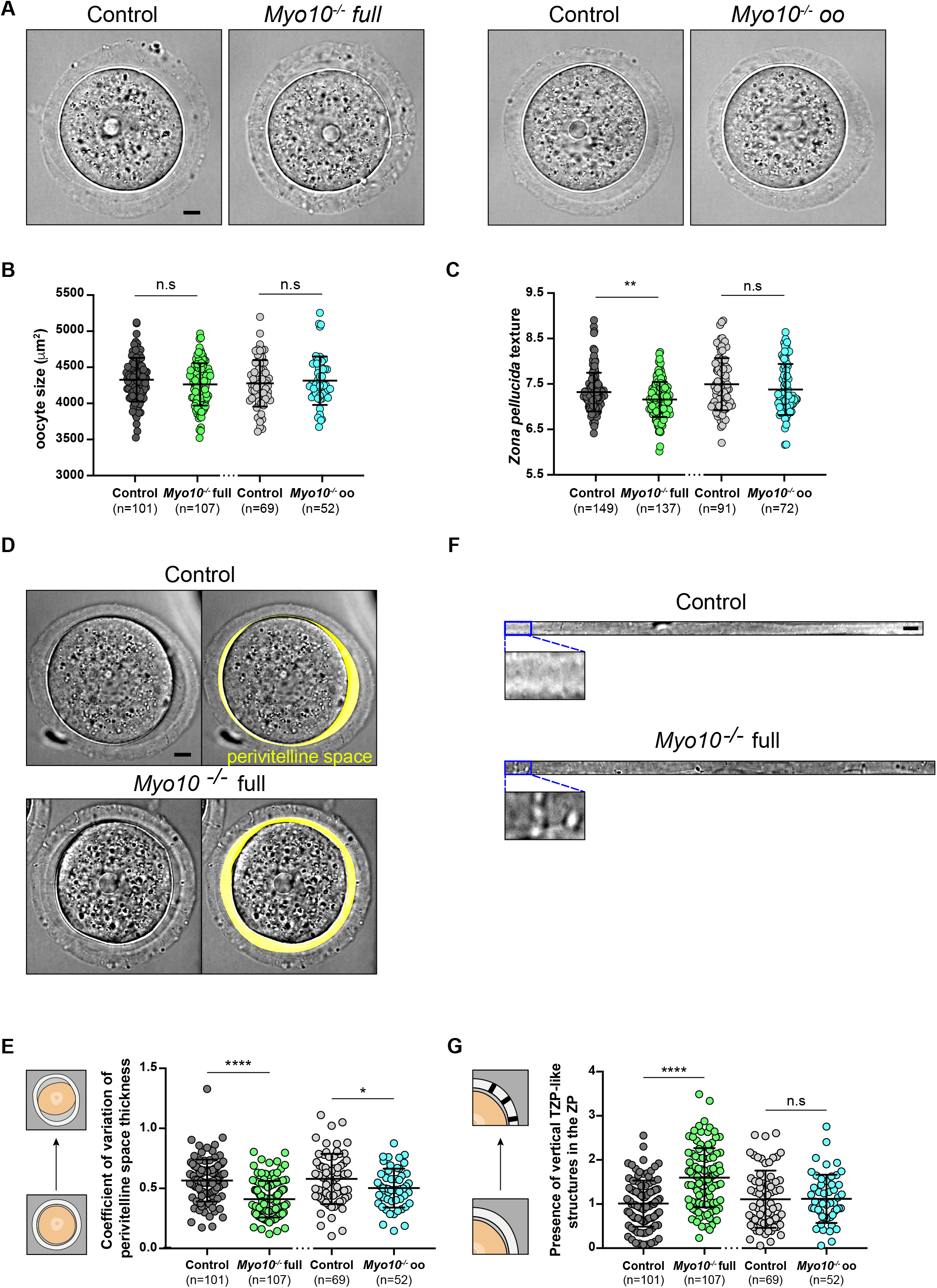
TZP-deprived oocytes have morphological alterations. **A)** Brightfield images of fully grown oocytes from the *Myo10* full knockout strain (left panel) and the *Myo10* oocyte knockout strain (right panel). For each panel, controls are on the left and *Myo10*^-/-^ oocytes on the right. Scale bar 10 μm. **B)** Scatter plot of the equatorial plane area of control and *Myo10*^-/-^ full oocytes (in dark gray and green, respectively) and control and *Myo10*^-/-^ oo oocytes (in light gray and blue, respectively). (n) is the number of oocytes analyzed. Data are mean±s.d. with individual data points plotted. Data are from four to thirteen independent experiments. Statistical significance of differences was assessed by a two-tailed unpaired t test, P value = 0.1243 (*Myo10*^-/-^ full oocytes compared to control oocytes) and by a two-tailed Mann Whitney test, P value = 0.7043 (*Myo10*^-/-^ oo oocytes compared to control oocytes). n.s not significant. **C)** Scatter plot of the *zona pellucida* texture of control and *Myo10*^-/-^ full oocytes (in dark gray and green, respectively) and control and *Myo10*^-/-^ oo oocytes (in light gray and blue, respectively). (n) is the number of oocytes analyzed. Data are mean±s.d. with individual data points plotted. Data are from four to thirteen independent experiments. Statistical significance of differences was assessed by a two-tailed Mann Whitney test, P value = 0.0029 (*Myo10*^-/-^ full oocytes compared to control oocytes) and by a two-tailed unpaired t test, P value = 0.1912 (*Myo10*^-/-^ oo oocytes compared to control oocytes). n.s not significant. **D)** Brightfield images of a fully grown control (upper panel) and *Myo10*^-/-^ full oocyte (bottom panel). For each panel, the right image displays the automatic segmentation of the oocyte perivitelline space (perivitelline space, yellow). Scale bar = 10 μm. **E)** Scatter plot of the coefficient of variation of the perivitelline space thickness, as represented by the schemes on the left. Control and *Myo10*^-/-^ full oocytes are in dark gray and green, respectively. Control and *Myo10*^-/-^ oo oocytes are in light gray and blue, respectively. (n) is the number of oocytes analyzed. Data are mean±s.d. with individual data points plotted. Data are from four to thirteen independent experiments. Statistical significance of differences was assessed by a two-tailed Mann Whitney test, P value < 0.0001 (*Myo10*^-/-^ full oocytes compared to control oocytes) and by a two-tailed unpaired t test, P value = 0.0308 (*Myo10*^-/-^ oo oocytes compared to control oocytes). **F)** Images of flattened *zona pellucida* obtained from oocyte segmentation. The top panel shows a control *zona pellucida* and the lower one a *Myo10*^-/-^ full *zona pellucida*. For each panel, the bottom image is a magnification of the brightfield image of the *zona pellucida* at the top. Scale bar 10 μm. **G)** Scatter plot of the presence of vertical TZP-like structures in the *zona pellucida* (ZP), as represented by the schemes on the left. Control and *Myo10*^-/-^ full oocytes are in dark gray and green, respectively. Control and *Myo10*^-/-^ oo oocytes are in light gray and blue, respectively. (n) is the number of oocytes analyzed. Data are mean±s.d. with individual data points plotted. Data are from four to thirteen independent experiments. Statistical significance of differences was assessed by a two-tailed unpaired t test with Welch’s correction, P value < 0.0001 (*Myo10*^-/-^ full oocytes compared to control oocytes) and by a two-tailed unpaired t test, P value = 0.9474 (*Myo10*^-/-^ oo oocytes compared to control oocytes). n.s not significant.

We further explored the differences in morphology between *Myo10*^-/-^ full and control fully grown oocytes using our machine-learning algorithm^22^ (Fig.2 D-G; S4 D-G), allowing us to detect in an unbiased manner the morphological features that differed the most between control and *Myo10*^-/-^ full oocytes. One of the most important feature used by our algorithm to distinguish control from *Myo10*^-/-^ full oocytes was the composition of the *zona pellucida* with the presence of vertical structures resembling TZPs crossing the *zona pellucida* (Fig.2 D, F, G; S4 D, G). Intriguingly, these structures were detected more frequently in *Myo10*^-/-^ full oocytes than in controls (Fig.2 G), despite the fact that total TZP density is decreased in *Myo10*^-/-^ full oocytes. The distribution of the perivitelline space around the oocyte was a second important feature used by our algorithm and was detected to be more uniform in *Myo10*^-/-^ full oocytes than in controls (Fig.2 D, quantification in E; S4 D, G), as if oocytes lacking TZPs were more homogeneously detached from the *zona pellucida*. Furthermore, our machine-learning algorithm identified *Myo10*^-/-^ full oocytes as more circular than control oocytes (Fig.S4 D, E, G), as indicated by a lower aspect ratio in TZP-deprived oocytes (Fig.S4 E). In accordance with this feature, the perimeter of *Myo10*^-/-^ full oocytes was slightly smaller than control oocytes for a similar area (Fig.S4 F).

Thus, using automatic approaches, we show that *Myo10*^-/-^ full oocytes can reach normal sizes but display morphological differences from controls, in particular regarding *zona pellucida* texture, a feature known to predict meiosis entry and maturation outcome^22^.

### Gene expression is modified in oocytes deprived of TZPs

We next wondered whether the differences both in the density of communicating structures between the oocyte and its follicular cells and in oocyte morphology translated into differences in oocyte developmental potential. The oocyte acquires its developmental potential during its growth, where it accumulates a large quantity of transcripts. Transcription decreases dramatically at the end of oocyte growth and increases after fertilization in the zygote^23^, thus protein synthesis during the meiotic divisions relies predominantly on the translational regulation of transcripts accumulated during growth, making this storage critical for oocyte quality^24–26^. Interestingly, follicular cells have been shown to modulate chromatin configuration and thus global transcriptional activity of the oocyte during its growth^24,27^. Since *Myo10*^-/-^full oocytes lack follicular cell contacts and potentially benefit less from exogenous transcriptional regulation, we tested whether their transcriptome differed from that of control oocytes.

By performing differential transcriptomic analysis by RNA-Seq between *Myo10*^+/-^ and *Myo10*^-/-^ full oocytes at the end of the growth phase^23^, we found 685 genes significantly (p-adj < 0.05) misregulated in *Myo10*^-/-^ full oocytes (Fig.3 A; Table.1), most of which were up-regulated (605 out of 685 representing 88.3% of deregulated transcripts). The majority of deregulated genes were protein coding (Fig.3 B), ten of which were validated by RT-qPCR (Fig.3 C). Among them, *Myo10* mRNA was confirmed to be down-regulated in *Myo10*^-/-^ full oocytes, validating both our RNA-Seq analysis and our genetic deletion tool (Fig.3 C). Since most of the deregulated transcripts in *Myo10*^-/-^ full oocytes were up-regulated, this could mean that the reduction in TZP density had resulted in either a higher transcription rate or increased stability of these transcripts.

**Figure 3:**
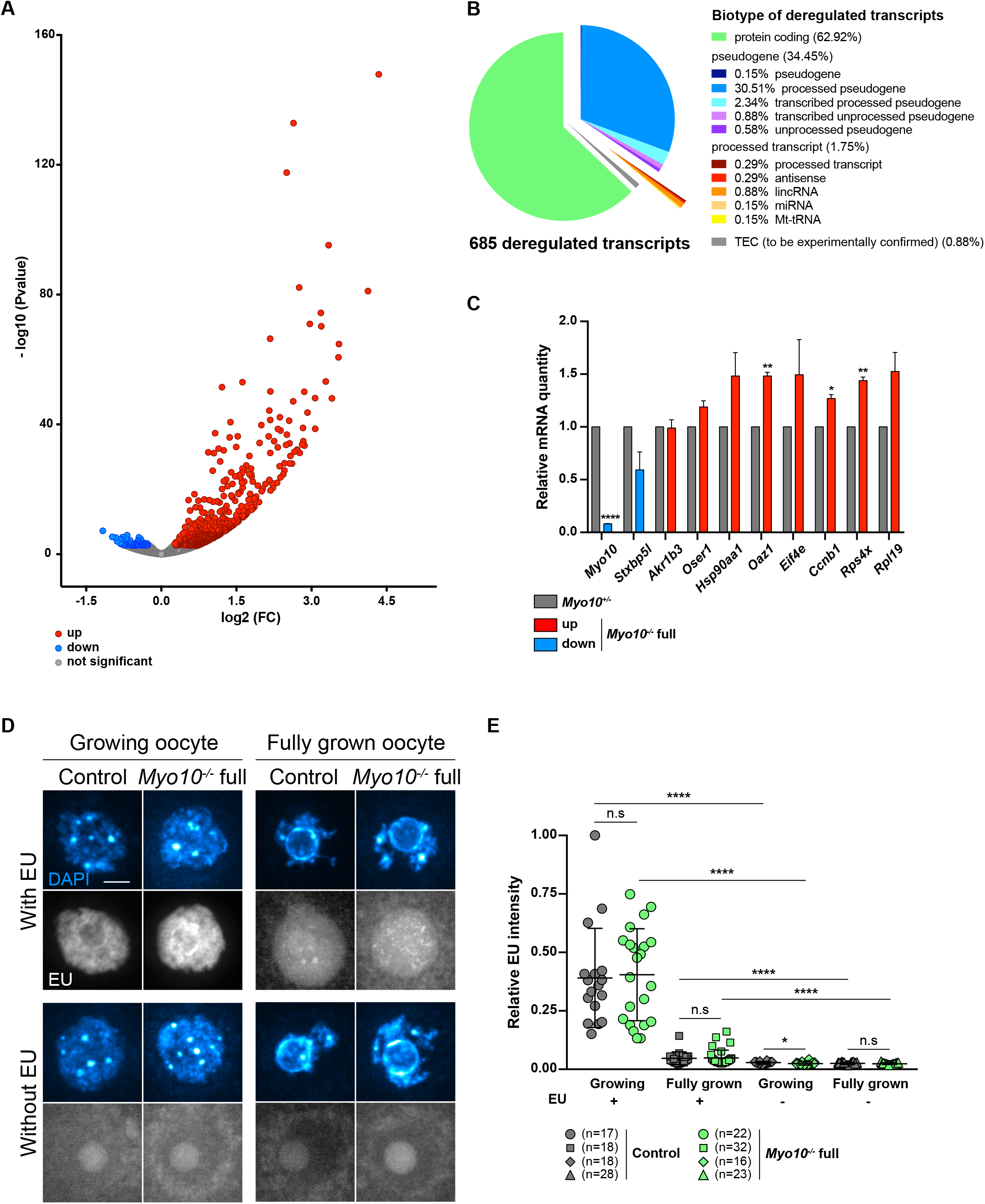
Gene expression is modified in TZP-deprived oocytes. **A)** Volcano plot of differential gene expression between fully grown *Myo10^+/-^* and *Myo10*^-/-^ full oocytes. Differential transcriptomic analysis was performed by RNA-Seq. Up-regulated transcripts are shown in red (up, 605 transcripts), down-regulated transcripts in blue (down, 80 transcripts, including *Myo10*), and non-deregulated transcripts in gray (not significant, 15,702 transcripts). Significance was set at P adj < 0.05. *Myo10*^-/-^ full oocytes were collected from two *Myo10^flox/flox^; Cre*^+^ mice and *Myo10^+/-^* oocytes from two *Myo10^wt/flox^; Cre*^+^ mice. For each condition, three biological replicates (each containing 10 oocytes) and three technical replicates were performed. **B)** Pie chart of deregulated transcript biotypes from the RNA-Seq analysis. lincRNA: long intergenic non-coding RNAs, miRNA: microRNA precursors, Mt-tRNA: transfer RNA located in the mitochondrial genome and TEC: To be Experimentally Confirmed. **C)** Bar graph of the relative mRNA quantity in fully grown *Myo10*^-/-^ full oocytes (colored bars) normalized to fully grown *Myo10^+/-^* oocytes (gray bars) performed by RT-qPCR. The selected mRNAs were chosen based on their biological relevance and their deregulatory strength (log2(FC) and P adj) in the RNA-Seq analysis described in A). mRNAs indicated as up- or down-regulated by RNA-Seq analysis are shown in red and blue, respectively. *Myo10*^-/-^full oocytes were collected from *Myo10^flox/flox^; Cre^-^* mice and *Myo10^+/-^* oocytes from *Myo10^wt/flox^; Cre*^+^ mice. For each condition, two biological replicates were performed (one containing 30 oocytes and the other 24 oocytes) from two different mice each time and two technical replicates were performed. Standard error of mean (SEM) is shown. For each mRNA, the mean of *Myo10^+/-^* oocytes was normalized to *Myo10^+/-^* oocytes. The mean and SEM for *Myo10^-/-^ full* oocytes *=* 0.08 ± 0.004 (*Myo10*); 0.59 ± 0.171 (*Stxbp5l*); 0.99 ± 0.079 (*Akr1b3*); 1.19 ± 0.059 (*Oser1*); 1.48 ± 0.222 (*Hsp90aa1*); 1.48 ± 0.034 (*Oaz1*); 1.49 ± 0.333 (*Eif4e*); 1.27 ± 0,037 (*Ccnb1*); 1.44 ± 0.034 (*Rps4x*); 1.53 ± 0.179 (*Rpl19*). Statistical significance of differences was assessed by two-tailed unpaired t tests. P value < 0.0001 (*Myo10*); *P value =* 0.1394 (*Stxbp5l*); 0.9001 (*Akr1b3*); 0.0864 (*Oser1*); 0.1612 (*Hsp90aa1*);0.0050 (*Oaz1*); 0.2765 (*Eif4e*); 0.0180 (*Ccnb1*); 0.0060 (*Rps4x*); 0.0987 (*Rpl19*). **D)** Sum of Z Projection images of oocytes stained with DAPI to label DNA (blue, top images of each panel) and incubated or not with 5-ethynyl uridine to label global RNA transcription (EU, bottom images of each panel). For each panel, control oocytes are on the left and *Myo10*^-/-^ full oocytes on the right. Growing and fully grown oocytes are on the left and right panels, respectively. Oocytes incubated with or without EU are in the upper and lower panels, respectively. For EU staining, contrast adjustment is similar between all conditions except for growing oocytes incubated with EU for which the signal intensity is too high to be shown with the same adjustment as the others. Scale bar 10 μm. **E)** Scatter plot of the relative EU signal intensity normalized to the DAPI signal intensity in control (dark gray) and *Myo10*^-/-^ full (green) growing or fully grown oocytes incubated or not with EU. (n) is the number of oocytes analyzed. Data are mean±s.d. with individual data points plotted. Data are from three to five independent experiments. Statistical significance of differences was assessed by two-tailed Mann Whitney tests, P value = 0.7688 (growing *Myo10*^-/-^ full oocytes with EU compared to growing control oocytes with EU), P value = 0.7564 (fully grown *Myo10*^-/-^ full oocytes with EU compared to fully grown control oocytes with EU), P value = 0.0326 (growing *Myo10*^-/-^ full oocytes without EU to growing control oocytes without EU), P value < 0.0001 (growing control oocytes with EU compared to growing control oocytes without EU), P value < 0.0001 (growing *Myo10*^-/-^ full oocytes with EU compared to growing *Myo10*^-/-^ full oocytes without EU), P value < 0.0001 (fully grown control oocytes with EU compared to fully grown control oocytes without EU) and P value < 0.0001 (fully grown *Myo10*^-/-^ full oocytes with EU compared to fully grown *Myo10*^-/-^ full oocytes without EU). Statistical significance of differences was assessed by a two-tailed unpaired t test to compare fully grown *Myo10*^-/-^ full oocytes without EU to fully grown control oocytes without EU, P value = 0.2647. n.s not significant.

We thus tested if this deregulation reflected a change in the global transcriptional rate of TZP-deprived oocytes^23^. Fully grown and smaller growing *Myo10*^-/-^ full oocytes were incubated with the uridine analog 5-ethynyl uridine (EU) to label RNA synthesis (Fig.3 D, E). Oocytes were also incubated without EU to serve as negative control. To measure global RNA synthesis, the signal intensity of EU was normalized to the DAPI signal intensity. Interestingly, global RNA synthesis was not different in growing and fully grown *Myo10*^-/-^ full oocytes compared to control oocytes (Fig.3 E).

In summary, we demonstrate here that lack of somatic contacts caused by decreased TZP density leads to a deregulation of oocyte gene expression at the end of its growth.

### TZP-deprived oocytes are prone to stop development in meiosis I despite correct spindle morphogenesis and chromosome alignment

We showed above that reduction of somatic contacts modulates the oocyte transcriptome. Importantly, oocytes depend on the transcripts synthesized during growth to develop until fertilization^25,26^. We thus tested whether TZP-deprived oocytes successfully underwent the developmental step following growth, namely meiotic divisions. After nuclear envelope breakdown (NEBD), the oocyte experiences an asymmetric division in size, leading to a large cell, the oocyte, and a tiny cell, the polar body. This size difference is essential to keep the nutrients stored during growth in the prospective fertilized oocyte for embryogenesis^28^. Importantly, we observed that *Myo10*^-/-^ full oocytes deprived of TZPs showed an increased frequency of developmental arrest at meiosis I, without extruding a polar body (Movie.S3; Fig.4 A, B, 38% of TZP-deprived oocytes do not extrude a polar body, compared to 15% in the controls), consistent with our morphological characterization using Oocytor (Fig.2 C). We previously showed that MYO10 expressed in oocytes is dispensable for meiotic divisions and female fertility^17^, suggesting that the meiotic arrest observed in a subset of *Myo10*^-/-^ full oocytes may be a consequence of decreased follicular cell contacts prior to meiosis resumption. We also observed that the subpopulation of *Myo10*^-/-^ full oocytes that extruded a polar body did so with a timing similar to controls (Movie.S3; Fig.4 A, C), indicating that complete loss of MYO10 and subsequent TZP depletion did not delay polar body extrusion in oocytes which could achieve it. Therefore, oocytes deprived of TZPs have a lower developmental potential than oocytes that retain a canonical density of follicular cell contacts, reinforcing the importance of the TZP-mediated dialogue between the oocyte and its niche for a fully efficient acquisition of oocyte competency.

**Figure 4:**
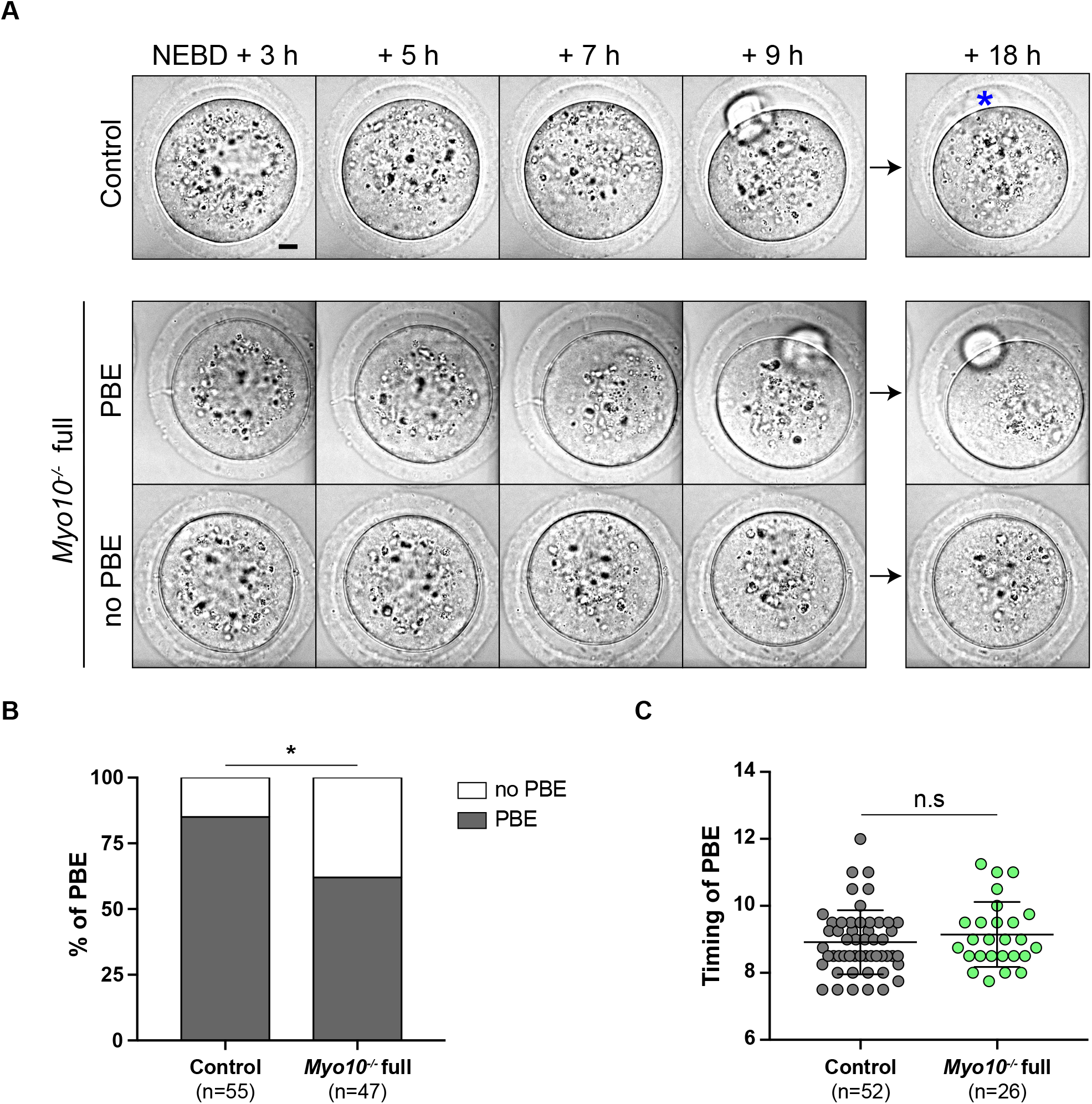
TZP-deprived oocytes tend to arrest in meiosis I. **A)** Brightfield images from spinning disk movies from 3 h to 18 h after NEBD. The top images show a control oocyte, the middle ones a *Myo10*^-/-^ full oocyte that extruded a polar body (PBE) and the bottom ones a *Myo10*^-/-^ full oocyte that did not extrude a polar body (no PBE). The blue asterisk indicates a lysed polar body. Scale bar 10 μm. **B)** Stacked bars of first polar body extrusion rate as a percentage of control and *Myo10*^-/-^ full oocytes. The gray bars represent the percentage of oocytes that extruded a polar body (PBE) and the white bars the percentage of oocytes that did not extrude a polar body (no PBE). (n) is the number of oocytes analyzed. Data are from three independent experiments. Statistical significance of differences was assessed by a two-sided Fisher’s exact test, P value = 0.0113. **C)** Scatter plot of the timing after NEBD of first polar body extrusion of control (dark gray) and *Myo10*^-/-^ (green) full oocytes. (n) is the number of oocytes analyzed. Data are mean±s.d. with individual data points plotted. Data are from six independent experiments. Statistical significance of differences was assessed by a two-tailed Mann Whitney test, P value = 0.3194. n.s not significant.

To gain insight into the origin of the meiotic arrest observed in some TZP-deprived oocytes, we first investigated whether the meiotic spindles were correctly formed in *Myo10*^-/-^ full oocytes. After NEBD, microtubules organize into a ball near the chromosomes that gradually bipolarizes to form a barrel-shape spindle^29–31^. To assess spindle morphogenesis, oocytes were stained with SiR-tubulin (SiR-tub) to label microtubules and followed throughout meiotic maturation. We observed that in both *Myo10*^-/-^ full oocytes that extruded a polar body and those that did not, meiosis I spindles bipolarized properly and within similar timings as in controls (Movie.S4; Fig.5 A-D). To further evaluate spindle morphogenesis in *Myo10*^-/-^ full oocytes, we measured spindle interpolar length and central width 7 hours after NEBD once spindle morphogenesis is complete and showed that *Myo10*^-/-^ full spindles were not significantly different from control spindles (Fig.5 B, D). Importantly, we observed that *Myo10*^-/-^ full oocytes that did not extrude a polar body arrested development in metaphase I, without experiencing anaphase (Movie.S4; Fig.5 A).

**Figure 5:**
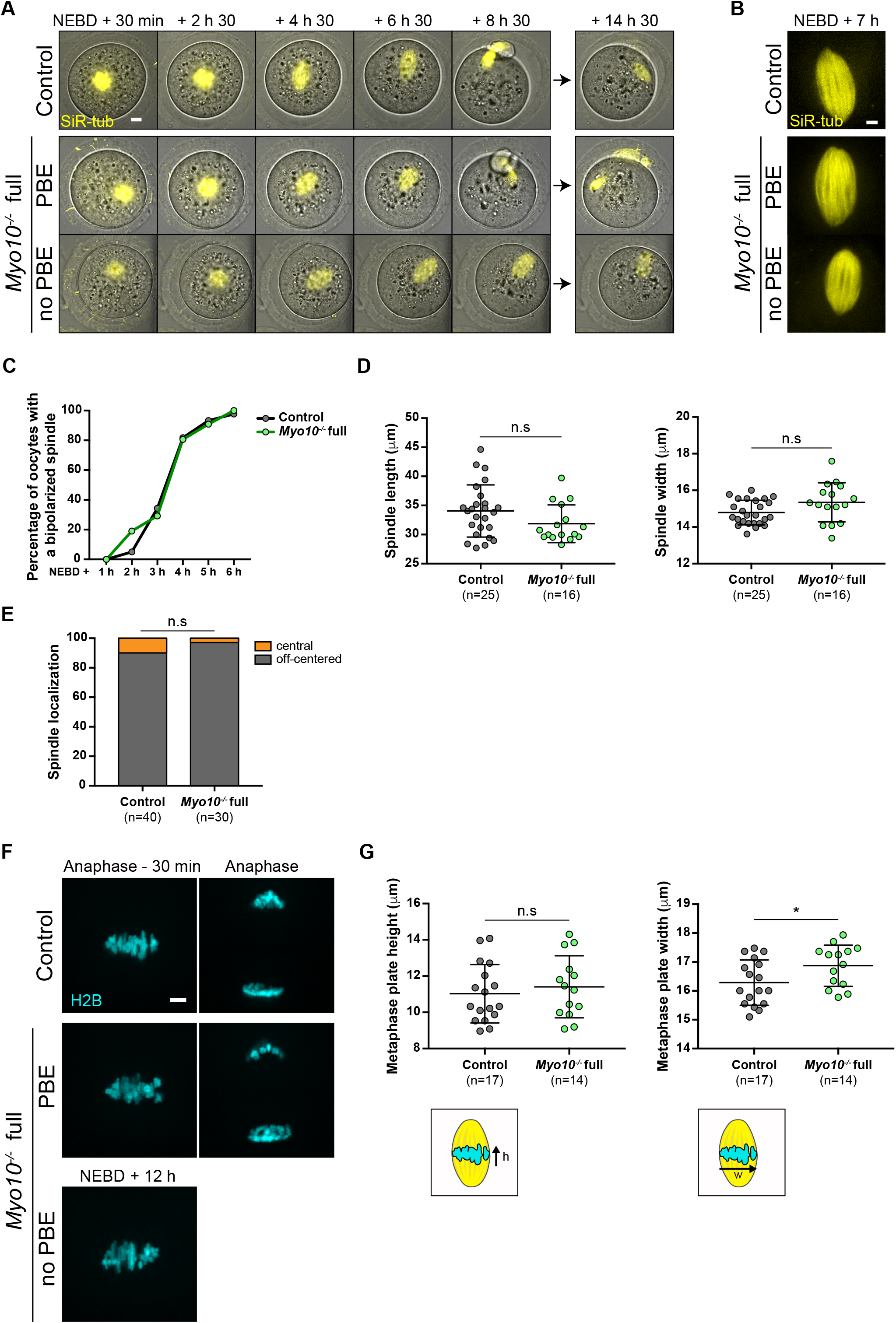
TZP-deprived oocytes have correctly formed and positioned spindles and aligned chromosomes. **A)** Max Intensity Z Projection images of oocytes stained with SiR-tubulin to label microtubules (SiR-tub, yellow) merged with corresponding brightfield images. Images are extracted from spinning disk movies from 30 min to 14 h 30 after NEBD. The top images show a control oocyte, the middle ones a *Myo10*^-/-^ full oocyte that extruded a polar body (PBE) and the bottom ones a *Myo10*^-/-^ full oocyte that did not extrude a polar body (no PBE). Scale bar 10 μm. **B)** Max Intensity Z Projection images of oocytes stained with SiR-tub (yellow) 7h after NEBD. The top image shows the spindle of a control oocyte, the middle one the spindle of a *Myo10*^-/-^ full oocyte that extruded a polar body (PBE) and the bottom one the spindle of a *Myo10*^-/-^ full oocyte that did not extrude a polar body (no PBE). Scale bar 5 μm. **C)** Graph of the percentage of control (dark gray) and *Myo10*^-/-^ (green) full oocytes with a bipolarized spindle as a function of time after NEBD. Data are from four independent experiments. For each time point, statistical significance of differences was assessed by a two-sided Fisher’s exact test. P value > 0.9999 for 1 h after NEBD; P value = 0.3433 for 2 h after NEBD; P value = 0.7765 for 3 h after NEBD; P value > 0.9999 for 4 h after NEBD; P value = 0.6937 for 5h after NEBD; P value > 0.9999 for 6h after NEBD. **D)** Scatter plots of spindle interpolar length (left) and central width (right) 7 h after NEBD in control (dark gray) and *Myo10*^-/-^ (green) full oocytes. (n) is the number of oocytes analyzed. Data are mean±s.d. with individual data points plotted. Data are from four independent experiments. Statistical significance of differences was assessed by a two-tailed unpaired t test, P value = 0.0989 for the length and by a two-tailed unpaired t test with Welch’s correction, P value = 0.0753 for the width. n.s not significant. **E)** Stacked bars of spindle cell localization 30 min before anaphase for oocytes that extruded a polar body or 12 h after NEBD for those that did not, as a percentage of control and *Myo10*^-/-^ full oocytes. Gray bars represent the percentage of oocytes with an off-centered spindle and orange bars the percentage of oocytes with a central spindle. (n) is the number of oocytes analyzed. Data are from four independent experiments. Statistical significance of differences was assessed by a two-sided Fisher’s exact test, P value = 0.3832. n.s not significant. **F)** Max Intensity Z Projection images from spinning disk movies of oocytes injected with H2B-GFP to label chromosomes (H2B, cyan) in a control oocyte (top images), a *Myo10*^-/-^ full oocyte that extruded a polar body (PBE, middle images) and a *Myo10*^-/-^full oocyte that did not extrude a polar body (no PBE, bottom image). Left images show chromosomes 30 min before anaphase for polar body extruding oocytes or 12h after NEBD for the *Myo10*^-/-^ full oocyte that did not extrude a polar body. Right images show chromosomes after anaphase. Scale bar 5 μm. **G)** Scatter plots of metaphase plate height (left) and width (right), as shown in the lower schemes, 30 min before anaphase for oocytes that extruded a polar body or 12 h after NEBD for those that did not. Control and *Myo10*^-/-^ full oocytes are in dark gray and green, respectively. (n) is the number of oocytes analyzed. Data are mean±s.d. with individual data points plotted. Data are from three independent experiments. Statistical significance of differences was assessed by two-tailed unpaired t tests. P value = 0.5331 for the height and P value = 0.0395 for the width. n.s not significant.

In addition to assessing spindle formation, we also examined meiosis I spindle positioning in *Myo10*^-/-^ full oocytes. During meiosis I, the spindle first assembles in the center of the oocyte and then migrates to the cell cortex by a process involving F-actin, resulting in asymmetric division in size^32–35^. We monitored the positioning of SiR-tub labeled spindles throughout meiosis I and analyzed their localization 30 minutes before anaphase for oocytes that extruded a polar body or up to 12 hours after NEBD for those that did not. The percentage of *Myo10*^-/-^ full oocytes with an off-centered spindle was not different from control oocytes (Movie.S4; Fig.5 A, E). In agreement with unaltered spindle positioning, we did not observe a defect in the organization of the Factin cytoplasmic network (including the actin cage) mediating spindle migration in fixed *Myo10*^-/-^ full oocytes labeled with phalloidin (Fig.S5 A-C). These results refine our understanding of the meiotic arrest of some *Myo10*^-/-^ full oocytes by showing that they arrest in metaphase I with a normally assembled and positioned spindle.

Since severe chromosome misalignment could prevent anaphase onset by sustaining the spindle assembly checkpoint in mouse oocytes^36^, we assessed chromosome alignment in *Myo10*^-/-^ full oocytes deprived of TZPs. Oocytes were injected with H2B-GFP to label chromosomes and chromosome alignment was monitored during meiosis I (Movie.S5; Fig.5 F, G). Interestingly, metaphase I chromosomes were aligned on the metaphase plate similarly to controls in *Myo10*^-/-^ full oocytes, as evidenced by a metaphase plate height not significantly different from controls 30 minutes before anaphase for oocytes that extruded a polar body or 12 hours after NEBD for those that did not (Fig.5 G). Therefore, the metaphase I arrest of some *Myo10*^-/-^ full oocytes is not caused by severe chromosome misalignment. Its origin remains to be further investigated.

### TZP-deprived oocytes produce fewer embryos, some cease development before the blastocyst stage, correlating with subfertility in females

Since TZP-deprived oocytes display oocyte-matrix defects, deregulated gene expression and a tendency to arrest in meiosis I, we sought to assess their developmental potential after fertilization. For that, we superovulated female littermates (*Myo10*^-/-^ *full* and control animals), mated them with male controls, and recovered embryos at E 0.5, *i.e*. at the zygote stage. First, we recovered more embryos for controls than for *Myo10*^-/-^ *full* animals for the same number of females (Fig.6 A compare the n, and compare the number of cells recovered from each female in the right images). Second, we recovered more unfertilized oocytes and more dead embryos in the *Myo10*^-/-^ full females compared to the controls (Fig.6 A black and gray bars respectively, and compare the number of 2-cell stage embryos at E1.5 for each female in the right images). Finally, we saw that among the embryos that survived, some stopped their development before reaching the blastocyst stage (Fig.6 B, black bars). All this suggests that the developmental potential of TZP-deprived oocytes is impaired. We therefore evaluated the fertility of *Myo10*^-/-^ *full* females by mating them with control males. The number of litters per couple, the mean number of pups per litter and thus the total number of pups per couple were significantly lower for *Myo10^-/-^ full* females than for controls (Fig.6 C), showing that *Myo10^-/-^ full* females are subfertile and arguing that TZP-mediated communication is important for the developmental potential of the oocyte and for female fertility.

**Figure 6:**
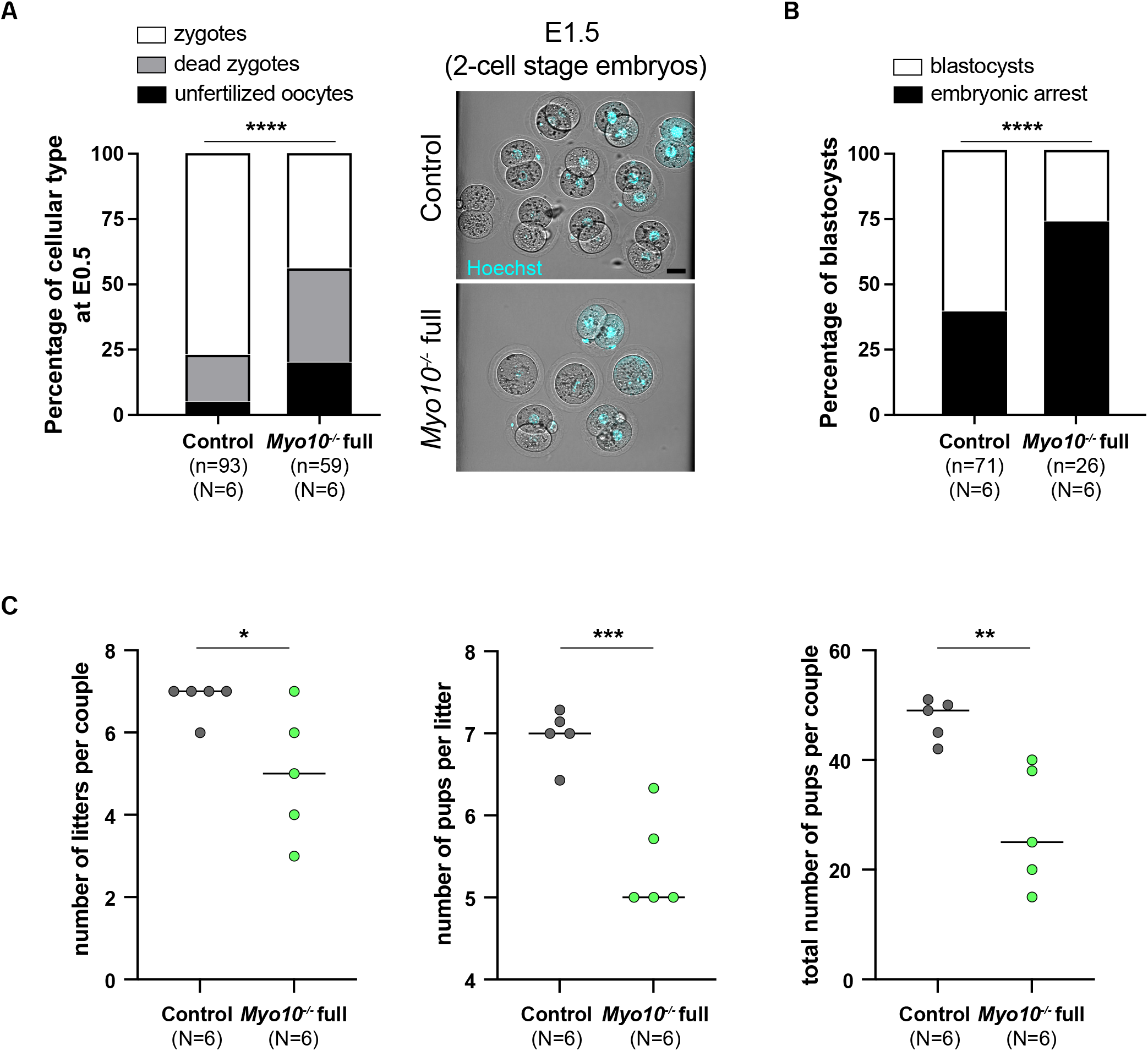
The development potential of TZP-deprived oocytes is altered, correlating with subfertility in females. **A)** On the left, stacked bars showing the percentage of unfertilized oocytes (black bars), dead zygotes (grey bars) and alive zygotes (white bars) recovered at E 0.5. (n) is the number of cells recovered, (N) the number of successful matings analyzed. Data are from three independent experiments. Statistical significance of differences was assessed with a Chi-square test, P value < 0.0001. On the right, cells recovered at E0.5 and cultured to E1.5 from a control female (upper panel) and a *Myo10*^-/-^ full female (lower panel), stained with Hoechst (blue) to visualize DNA. Scale bar 40 μm. **B)** Stacked bars showing the percentage of embryonic arrest (black bars) and blastocysts (white bars). (n) is the number of cells recovered, (N) the number of successful matings analyzed. Data are from three independent experiments. Statistical significance of differences was assessed by a two-sided Fisher’s exact test, P value < 0.0001. **C)** Scatter-plots of the number of litters per couple, number of pups per litter and total number of pups per couple from *Myo10^wt/wt^; Cre*^+^ female mice (control, grey) and *Myo10^flox/flox^; Cre*^+^ female mice (carrying *Myo10*^-/-^ full oocytes, green). Six independent matings (N) were set for six months each with *Myo10^wt/wt^; Cre^-^* males. The statistical significance of the differences were assessed with two-tailed unpaired t tests, P value = 0.04 (for the number of litters per couple), P value = 0.0009 (for the number of pups per litter) and P value = 0.0052 (for the total number of pups per couple).

## Discussion

In this study, we aimed to gain more insight into the contribution of cellular protrusions established by surrounding somatic cells to postnatal oocyte development. By genetically deleting the transzonal projection component MYO10 in all murine cell types, including follicular cells, we greatly reduced the density of oocyte-follicular cell contacts and assessed the resulting oocyte development.

Surprisingly, although complete loss of MYO10 decreased by fivefold the density of TZPs in fully grown oocytes, *Myo10*^-/-^ full oocytes grew to normal sizes (Fig.2 A, B), and were arrested in prophase. Hence, and in contrast to previous studies where the impairment of oocyte-follicular cell communication by deletion of connexin 37 or GDF9 led to oocytes respectively smaller or larger than controls^18–20^, reducing TZP density did not prevent oocytes from growing to a canonical size. Interestingly, a small fraction of TZPs remained in *Myo10*^-/-^ full oocytes, implying that a subpopulation of TZPs may be MYO10 independent, and potentially explaining how oocytes managed to grow to canonical size and maintained the meiotic arrest in prophase despite likely reduced intercellular communication.

Even though TZP-deprived oocytes grew to normal sizes, we identified morphological differences, mostly related to the integrity of the *zona pellucida*. The *zona pellucida*, which is the extracellular matrix surrounding the oocyte, is composed of proteins secreted by the oocyte throughout growth and assembled in cross-linked filaments^37,38^. Using our machine-learning algorithm to phenotype oocytes, we found that the *zona pellucida* was more homogeneously detached in TZP-deprived oocytes. It is likely that reduction in TZP density, and consequently the decrease in adherens junctions between the oocyte and follicular cells^9,11^, led to this apparent looser oocyte-matrix adhesion. Furthermore, our machine-learning algorithm detected TZP-like structures more readily in *Myo10*^-/-^ full oocytes than in controls, despite their decrease in TZP density. This could reflect an alteration in the structure of the remaining TZPs, or alternatively a difference in the structure of the *Myo10*^-/-^ full *zona pellucida* that would make the remaining protrusions crossing it more apparent. All these features taken together may reflect impaired assembly or maintenance of the *zona pellucida* in oocytes with reduced cell-cell contacts, since the *zona pellucida* is a viscoelastic extracellular matrix whose properties could change as a function of the crosslinking generated by the number of TZPs passing through it^37–38^. In support of this, we occasionally observed some follicular-like cells located ectopically within the periviteline space of TZP-deprived oocytes, a phenotype not observed in control oocytes (Fig.S4 C, see red arrow). The same ectopic localization was observed in GDF9-deficient mice, which also displayed oocytes with reduced TZP density, and such abnormal localization of follicular cells was correlated with low oocyte viability^20^. Interestingly, the presence of follicular-like cells inside the perivitelline space has been observed in mice lacking ZP1^39^, the secreted *zona pellucida* protein that mediates the cross-linking of the matrix filaments^37–38^. By modulating matrix filament cross-linking, TZPs may therefore be involved in preserving *zona pellucida* integrity, which is important for fertilization^37–38^. Interestingly, we recovered more unfertilized oocytes from *Myo10*^-/-^ *full* females compared to controls after mating, which suggests a potential effect of the observed alteration of the *zona pellucida* on fertilization (Fig.6 A).

Importantly, TZP-deprived oocytes have lower developmental competence than control oocytes, as they more frequently arrest at metaphase of the first meiotic division, and produce fewer embryos with lower viability, impairing female fertility. Consistent with our results, a decrease in oocyte developmental potential correlated with reduced TZP density was also observed in oocytes lacking the focal adhesion kinase Proline-rich tyrosine kinase 2^40^. In *Myo10*^-/-^ full oocytes, the basis of the meiotic arrest observed in a subpopulation of oocytes remains to be deciphered. This developmental arrest may be the readout of the altered oocyte transcriptome since we revealed by RNA-Seq analysis that gene expression prior to meiosis resumption is modified in TZP-deprived oocytes. As oocytes undergo meiotic divisions relying on transcripts synthesized and stored during growth^25–26^, an impairment in this storage could compromise oocyte quality. Interestingly, some deregulated transcripts in TZP-deprived oocytes, including the most deregulated ones, are involved in the oxidative stress response, the metabolism of polyamine and cell cycle transitions (*Oser1, Akr1b3, Oaz1, Psma5, Cyclin B1*, Fig.3 C, Table 1), pathways that when deregulated are known to affect subsequent meiotic maturation and early embryogenesis^41–46^.

In conclusion, we have shown that cytoplasmic protrusions of surrounding somatic cells enhance the developmental potential of the oocyte. Consistent with others studies, we suggest that TZPs contribute to zona pellucida integrity and oocyte-matrix adhesion - directly as an intercellular contact structure or indirectly through metabolite exchange optimizing oocyte quality and the establishment of a robust zona pellucida - thus playing a role in the structural integrity of the ovarian follicle. In addition, we propose a novel role for TZPs as modulators of the synthesis/stability of specific oocyte transcripts at the end of growth that are required for successful meiotic divisions and early embryonic development. Interestingly, as the density of TZPs decreases with maternal age^7^, investigating their contribution to oocyte quality could be of great benefit to further unraveling female fertility, which decreases with maternal age. This may be of particular interest to increase the efficiency of assisted reproductive technologies. More broadly, our results may be valuable for the comprehension of other cellular models relying on distant cell-cell contact communication to function^47^. It may notably provide clues as to how cellular development is modulated by exogenous regulation of gene expression through direct intercellular dialogue.

## Acknowledgments

We thank the Verlhac/Terret lab for comments on the manuscript and discussions; C. Antoniewski and N. Naouar from the IBPS ARTbio bioinformatics platform for their services and discussion. The STED microscopy was performed at the Orion Platform (member of France–Bioimaging ANR-10-INBS-XX) of the Center for Interdisciplinary Research in Biology (UMR7241/U1050) of Collège de France. The OMX microscopy was performed at the Cell and Tissue Imaging Platform -PICT-IBiSA (member of France–Bioimaging ANR-10-INBS-04) of the Genetics and Developmental Biology Department (UMR3215/U934) of Institut Curie, supported by the European Research Council (ERC EPIGENETIX N°250367). The work at IBENS Genomic Platform was supported by the France Génomique national infrastructure, funded as part of the “Investissements d’Avenir” program managed by the Agence Nationale de la Recherche (reference: ANR-10-INBS-09). This work was supported by the Fondation pour la Recherche Médicale (FRM Label EQU201903007796 to MHV), from the Agence Nationale de la Recherche (ANR-18-CE13 to MHV and Auguste Genovesio, IBENS/ENS, ANR-16-CE13 to MET) and from the France Canada Research Fund (FCRF 2017 to MET and Hugh J Clarke, McGill University). FC obtained a grant from the Fondation pour la Recherche Médicale for her 4^th^ PhD year (FDT202001010906). This work has received support from the Fondation Bettencourt Schueller, support under the program « Investissements d’Avenir » launched by the French Government and implemented by the Agence Nationale de la Recherche, with the references: ANR-10-LABX-54 MEMO LIFE, ANR-11-IDEX-0001-02 PSL* Research University.

## Author contributions

The project was conceived by F.Cr., M.H.V. and M.E.T and supervised by M.E.T. Additional input was provided by H.J.C. M.H.V. was the first to observe the decrease in TZP density. All experiments on mouse oocytes were performed by F.Cr. with assistance from A.E. and M.E.T when needed. F.Cr. and M.E.T analyzed and quantified most experiments. G.L. performed the machine-learning approach to phenotype oocytes. C.D.S. performed and analyzed RT-qPCR experiments. A.F.T and J.L.M supervised and performed all experiments on mouse embryos. M.B and A.C supervised and performed ovary transparization and data acquisition. J.D, T.P and A.D performed super-resolution acquisitions (STED and OMX). F.Co did the cDNA libraries and performed the RNA-seq, which was analyzed by the IBPS bioinformatics facility (ARTbio). F.Cr wrote the original draft of the manuscript, which was reviewed and corrected by M.E.T and edited by all authors.

## Material and methods

### Mouse strain generation and genotyping

To generate *Myo10^wt/flox^* mice, two *loxP* sites flanking *Myo10* exons 23-25 and an FRT-flanked neomycin selection cassette located between *Myo10* exons 22 and 23 were inserted in the genome of C57BL/6 mice by genOway. Mice were then mated with *Zp3-Cre* mice already housed in the laboratory.

To obtain *Myo10* full knockout mice (*Myo10^-/-^* full), the neomycin selection cassette was retained to allow its promoter to interfere with *Myo10* expression independently of Cre expression^48^ (Fig.S1 A). We used the following primers to detect the presence of the neomycin selection cassette at the *Myo10* locus: 5′-ACA GCC CAT ATC ACT GTC TAG AGA CCC ATT-3′, 5′-ATC GCC TTC TAT CGC CTT GAC G-3′ and 5′-GAG GAT CCA GAC TTG GAC CCG GTC-3′.

Mice conditionally deleted for *Myo10* in oocytes (*Myo10^-/-^* oo mice) were previously generated from the above strain by removing the neomycin selection cassette after mating with *FLPo* mice^17^ (Fig.S1A). To detect *Cre*-mediated excision at the *Myo10* locus after removing of the neomycin selection cassette, the following PCR primers were used: for the *Cre*, 5′-GCG GTC TGG CAG TAA AAA CTA TC-3′ and 5′-GTG AAA CAG CAT TGC TGT CAC TT-3′; and for the *Myo10*, 5′-ACC CCA GTA CTT GTT CAT ACA TCC TAT ATC CTA CA-3′, 5′-GAC TAC ACC ATT CTG AAT GTG CCT GAT CTC-3′ and 5′-GAG TAT CTG CCA TCT TGT CCC TAA AGG TGG-3′.

For the *Myo10* full knockout strain, we used *Myo10^wt/wt^; Cre^+^, Myo10^wt/flox^; Cre^+^, Myo10^wt/D^; Cre^+^, Myo10^wt/flox^; Cre^-^* as control mice and *Myo10^flox/flox^; Cre^+^, Myo10^flox/D^; Cre^+^, Myo10^flox/flox^; Cre^-^* as *Myo10*^-/-^ full mice. For the *Myo10* oocyte knockout strain, we used *Myo10^wt/flox^; Cre^+^, Myo10^wt/flox^; Cre^-^, Myo10^flox/flox^; Cre^-^* as control mice and *Myo10^flox/flox^; Cre*^+^ as *Myo10*^-/-^ oo mice.

### Immunoblotting

Immunoblotting was performed as in Ref.15 on extracts from *Myo10^wt/flox^; Cre^+^, Myo10^flox/flox^; Cre*^+^ and *Myo10^flox/flox^; Cre^-^* mice from the *Myo10* full knockout strain. Whole liver, whole spleen and one kidney per mouse were extracted, weighed and immediately added to 1X Laemmli sample buffer (50 mM Tris, pH 7.4, 2% SDS, 6% glycerol, 0.1 M DTT and 0.25% bromophenol blue) at 1 mL/20 mg wet tissue weight and boiled for 15 min. Samples were crushed with a glass potter to dissociate the tissue and the lysates were centrifuged at 20,000 rpm for 30 min at room temperature. The pellets were discarded and the supernatants were stored at −80°C. Final loading volumes were adjusted for immunoblot based on quantification of the actin band from preliminary tests, so that the same protein amount is deposited in all the wells. Samples were boiled 1 min and subjected to SDS-PAGE using a 4-15% Mini-PROTEAN TGX Stain-Free Protein Gel (Bio-Rad Ref. 4568085). After SDS-PAGE, lysates were transferred to Immobilon-P PVDF Membrane (Sigma-Aldrich Ref. IPVH00010) at room temperature in transfer buffer (25 mM Tris, 192 mM Glycine, 0.3% SDS and 20% methanol) using a Trans-Blot SD Semi-Dry Transfer Cell (Bio-Rad) set at 100 mA constant power with a volt limit of 25 V for 45 min. Prior to transfer, the PVDF membrane was wetted in methanol, rinsed in distillated water and equilibrated 10 min in transfer buffer. After the transfer, the membrane was washed in 50 mM Tris, 150 mM NaCl, 0.05% Tween 20 (TBST thereafter) and blocked for 1h in TBST containing 5% dried milk powder. The membrane was washed in TBST and incubated for 1h in TBST containing 4% dried milk and rabbit anti-MYO10 (1:300, Sigma-Aldrich Ref. HPA024223) as primary antibodies. The membrane was washed in TBST, blocked for 1h in TBST containing 5% dried milk powder, rinsed in TBST and incubated for 45 min in TBST containing 4% dried milk and donkey anti-rabbit IgG HRP-linked (1:10000, Amersham Ref. NA934) as secondary antibodies. The membrane was washed in TBST and incubated 5 min in SuperSignal West Femto Maximum Sensitivity Substrate (Thermo Fisher Scientific Ref. 34095). Protein expression was revealed using a chemiluminescence Image Analyzer (FUSION FX; Vilber, 20 sec illumination).

The membrane was then rinsed in TBST and stripped 10 min in Restore Western Blot Stripping Buffer (Life Technology Ref. 21059). The membrane was washed in TBST and incubated for 45 min in TBST containing 4% dried milk and HRP mouse anti-beta Actin AC-15 (1:25000, abcam Ref. ab49900). The membrane was washed in TBST and incubated 1 min in ECL Western Blotting Substrate (Thermo Fisher Scientific Ref. 32109). Protein expression was revealed using a chemiluminescence Image Analyzer (FUSION FX; Vilber, 20 sec illumination).

### Whole ovary immunostaining and clearing

Ovaries collected from two control mice (*Myo10^wt/flox^; Cre*^+^, 9 and 12 weeks old) and two *Myo10*^-/-^ full mice (*Myo10^flox/flox^; Cre*^+^, 9 and 12 weeks old) were fixed in 1X PBS, 4% paraformaldehyde for 4h with agitation at room temperature and subsequently washed in 1X PBS where they were stored at 4°C. Bleaching, immunostaining and clearing were performed as in Ref.21. Ovaries were first dehydrated in increasing concentrations of methanol (1X PBS, methanol 50%, 80% and 100% methanol twice) for 1h each time with agitation at room temperature. Ovaries were then bleached in 100% methanol + 6% hydrogen peroxide overnight at 4°C and rehydrated in decreasing concentrations of methanol (100% twice, 1X PBS, methanol 80% and 50%) for 1h each time with agitation at room temperature. Ovaries were blocked and permeabilized in 1X PBS, 0.2% gelatin, 0.5% Triton X-100 (PBSGT thereafter) for 4 days with rotation at room temperature. Antibody incubations were performed in PBSGT + 10X saponin (10 mg/mL) at 37°C with rotation at 70 rpm for one week for primary antibodies and overnight for secondary antibodies. Washes between antibody incubations and after secondary antibody incubation were done in PBSGT for 1day with rotation at room temperature. We used rabbit anti-MYO10 (1:1000, Sigma-Aldrich Ref. HPA024223) and rat anti-MECA-32 (1:500, BD Pharmingen Ref. 550563, data not shown) as primary antibodies and Cy3-conjugated donkey anti-rabbit IgG (1:500, Jackson Immuno research Ref. 711-165-152) and Alexa Fluor 790-conjugated donkey anti-rat IgG (1:250, Jackson Immuno research Ref. 712-655-153) as secondary antibodies (data not shown). Incubation with TO-PRO-3 (1:100, ThermoFisher Scientific Ref. T3605) was performed along with secondary antibody incubation to stain nucleic acid. Ovaries were embedded in 1X TAE, 1.5 % agarose and cleared using an adapted 3DISCO clearing protocol with rotation at 12 rpm and protected from light at room temperature. Ovaries were dehydrated in H20, 50% tetrahydrofuran (THF anhydrous with 250 ppm butylated hydroxytoluene inhibitor, Sigma-Aldrich Ref.186562) overnight and in H20, 80% THF and 100% THF twice for 1 h 30 each time. Ovaries were then incubated in dichloromethane (DCM, Sigma-Aldrich Ref.270997) for 30 min for delipidation and cleared overnight in benzyl ether (DBE, Sigma-Aldrich Ref.108014).

### Follicle/oocyte collection and culture

Preantral follicles, cumulus-oocyte complexes and oocytes isolated from follicular cells were collected from 2- to 5-month-old mice in M2 + BSA medium supplemented with 1μM milrinone^49^ to maintain oocytes arrested in prophase I. Oocytes were released from follicular cells by repeated pipette aspirations. Meiosis resumption was induced by transferring oocytes to M2 + BSA medium free of milrinone. Culture and live imaging were performed at 37°C under oil. To assess the rate of polar body extrusion, oocytes were cultured overnight in M2 + BSA medium in a Heracell 150i CO_2_ Incubator (Thermo Fisher Scientific) and analyzed the next day for polar body extrusion.

### Embryo recovery and culture

Embryos were isolated from superovulated female mice (*Myo10^flox/flox^; Cre^+^, Myo10^flox/flox^; Cre^-^* as *Myo10*^-/-^ full mice and *Myo10^wt/wt^; Cre^+^, Myo10^wt/wt^; Cre^-^* as control mice) mated with male mice (*Myo10^wt/wt^; Cre*^-^). Superovulation of female mice was induced by intraperitoneal injection of 5 international units (IU) pregnant mare’s serum gonadotropin (Ceva, Syncro-part), followed by intraperitoneal injection of 5 IU human chorionic gonadotropin (MSD Animal Health, Chorulon) 44–48 h later. Embryos were recovered at E 0.5 from plugged females by opening the ampulla followed by a brief treatment with 37°C 0.3 mg/mL hyaluronidase (Sigma, H4272-30MG) and washing in 37°C FHM. When ampulla was not present, embryos were recovered by flushing oviducts with 37°C FHM (Millipore, MR-122-D) using a modified syringe (Acufirm, 1400 LL 23). Embryos were handled using an aspirator tube (Sigma, A5177-5EA) equipped with a glass pipette pulled from glass micropipettes (Blaubrand intraMark or Warner Instruments). Embryos were placed in KSOM (Millipore, MR-107-D) supplemented with 0.1% BSA (Sigma, A3311) in 10 mL droplets covered in mineral oil (Sigma, M8410). Embryos were cultured in an incubator under a humidified atmosphere supplemented with 5% CO_2_ at 37°C for 5 days. Embryos were scored for survival and embryonic stage from E0.5 to E4.5.

### Fluorescent probes for live imaging

Oocytes were incubated for 1 h in M2 + BSA, milrinone medium supplemented with 100 nM SiR-actin (Spirochrome Ref. SC006) to label F-actin, 5 μg/mL FM 1-43 (Thermo Fisher Scientific Ref. T35356) to label membranes or 100 nM SiR-DNA (Spirochrome Ref. SC007) to label DNA. Oocytes were incubated in M2 + BSA medium supplemented with 100 nM SiR-tubulin (Spirochrome Ref. SC006) at least 1 h before the start of spinning disk movies and throughout movie acquisitions to label microtubules. Embryos at E1.5 were incubated 15 minutes in M2 + BSA medium supplemented with 5 ng/mL Hoechst (Invitrogen Ref. H3570) to label DNA.

### *In vitro* cRNA synthesis and oocyte microinjection

The pRN3-Histone(H2B)-GFP plasmid^50^ was linearized with SfiI restriction enzyme and *in vitro* transcribed using the mMESSAGE mMACHINE T3 Transcription Kit (Ambion Ref. AM1348). cRNAs were purified using the RNeasy Mini Kit (QIAGEN Ref. 74104), centrifuged at 4°C and microinjected into prophase I arrested oocytes with a FemtoJet microinjector (Eppendorf). Oocytes were maintained in prophase I for 2 h to express cRNAs before meiosis resumption.

### Immunostaining and global transcription detection

Preantral follicles, cumulus-oocyte complexes and oocytes isolated from follicular cells were immunostained as in Ref.17. Incorporation and detection of 5-ethynyl uridine (EU) into nascent oocyte RNAs were performed similarly to Ref.23. For all protocols, follicles, complexes and oocytes were washed in M2 + PVP medium and adhered to gelatin and polylysine-coated coverslips before fixation^51^.

#### Phalloidin labeling of transzonal projections

Preantral follicles and cumulus-oocyte complexes were fixed and permeabilized in 1X PBS, HEPES (100 mM, pH 7), EGTA (50 mM, pH 7), MgSO_4_ (10 mM) buffer with 2% Triton X-100 and 0.3% formaldehyde for 30 min at 37°C. Follicles and complexes were then washed in 1X PBS where they were stored overnight or 2 days at 4°C. Follicles and complexes were further permeabilized in 1X PBS, 0.5% Triton X-100 for 10 min and washed in 1X PBS and 1X PBS, 0.1% TWEEN 20 (PBSTw thereafter). Blocking and antibody incubations were performed in PBSTw, 3% BSA at room temperature for 30 min for blocking, 1 h 30 for primary antibody incubation and 1h for secondary antibody incubation. Washes between antibody incubations and after secondary antibody incubation were done in PBSTw at room temperature and an additional wash in 1X PBS was performed before mounting. For spinning disk and OMX microscopy, follicles and complexes were mounted on slide wells filled with VECTASHIELD Antifade Mounting Medium with or without DAPI (Vector Laboratories Ref. H-1200 and H-1000). We used rabbit anti-MYO10 (1:200, Sigma-Aldrich Ref. HPA024223) as primary antibodies and Cy3-conjugated donkey anti-rabbit IgG (1:150, Jackson ImmunoResearch Ref. 711-165-152) as secondary antibodies. Incubation with Alexa Fluor 488-conjugated Phalloidin (10 U/mL, Thermo Fisher Scientific Ref. A12379) was performed concurrently with secondary antibody incubation to stain F-actin. For STED microscopy, cumulus-oocyte complexes and oocytes freed from follicular cells were mounted on slide wells filled with Abberior liquid antifade mounting medium (Ref. MM-2009). We used rabbit anti-MYO10 (1:200, Sigma-Aldrich Ref. HPA024223) as primary antibodies and Abberior STAR 580-conjugated anti-rabbit (1:200, Ref. ST580-1002) as secondary antibodies. Incubation of Abberior STAR RED-conjugated Phalloidin (1:200, Ref. STRED-0100) was performed with secondary antibody incubation to stain F-actin.

#### Phalloidin labeling of oocyte cytoplasmic F-actin

Oocytes isolated from antral follicles were cleared of the *zona pellucida* in acid Tyrode 1 h – 1 h 30 before fixation. Immunostaining was performed using a modified protocol from Ref.33. Oocytes were fixed and permeabilized 6 h after NEBD in 1X PBS, HEPES (100 mM, pH 7), EGTA (50 mM, pH 7), MgSO4 (10 mM) buffer with 0.2% Triton X-100 and 2% formaldehyde for 30 min at 37°C. Oocytes were then washed in 1X PBS, 0.1% Triton X-100 (PBSTr thereafter) where they were stored overnight at 4°C. The remaining immunostaining protocol is similar to that for phalloidin TZP labeling for spinning disk and OMX microscopy, except that oocytes were directly blocked the day after fixation and blocking and antibody incubations were performed in PBSTr, 3% BSA. Oocytes were mounted on slide wells filled with VECTASHIELD Antifade Mounting Medium with DAPI (Vector Laboratories Ref. H-1200). We used mouse anti-α-Tubulin (1:200, Sigma-Aldrich Ref. T8203) as primary antibodies and Cy3-conjugated donkey anti-mouse IgG (1:150, Jackson ImmunoResearch Ref. 715-165-151) as secondary antibodies. Incubation with Alexa Fluor 488-conjugated Phalloidin (10 U/mL, Thermo Fisher Scientific Ref. A12379) was performed along with secondary antibody incubation.

#### EU labeling of nascent oocyte RNAs

As in Ref.23, oocytes arrested in prophase I isolated from follicular cells were cleared of the *zona pellucida* in M2 + BSA, milrinone medium with 0.4% pronase. EU incorporation and detection were performed using the Click-iT RNA Alexa Fluor 488 Imaging Kit (Thermo Fisher Scientific Ref. C10329). Oocytes were incubated in M2 + BSA, milrinone medium with or without 0.5 mM EU for 3 h at 37°C and fixed in 1X PBS, 4% paraformaldehyde for 30 min at 37°C. Oocytes were subsequently washed in 1X PBS where they were stored overnight at 4°C. Oocyte permeabilization and washes were done at room temperature in 1X PBS, 0.5% Triton X-100 for 10 min and in 1X PBS, respectively. EU detection was performed as indicated by the manufacturer and oocytes were then washed twice in Click-iT reaction rinse buffer and mounted on slide wells filled with VECTASHIELD Antifade Mounting Medium with DAPI (Vector Laboratories Ref. H-1200).

### Imaging

#### Spinning disk microscopy

Images were acquired with a Plan-APO 40×/1.25 NA objective on a Leica DMI6000B microscope enclosed in a thermostatic chamber (Life Imaging Service) equipped with a Retiga 3 CCD camera (QImaging) coupled to a Sutter filter wheel (Roper Scientific) and a Yokogawa CSU-X1-M1 spinning disk. Data were collected with Metamorph software (Universal Imaging).

#### STED microscopy

STED imaging was performed with a STEDYCON module from Abberior mounted on a Zeiss AxioObserver 7 inverted-Videomicroscope coupled to a sCMOS camera (Hamamatsu Flash 4) and a PlanAPO 100× oil-immersion objective (NA = 1.46) for an effective pixel size of 50 nm (Zeiss).

#### OMX microscopy

structured illumination (3D-SIM) was performed using a DeltaVision OMX Blaze microscope (GE Healthcare) coupled to a Photometrics Evolve 512 EMMCD camera (Photometrics, Tucson, AZ) and a PlanAPO 60× oil-immersion objective (NA =1.4) for an effective pixel size of 80 nm and 40 nm after reconstruction (Olympus, Melville, NY).

#### Light-sheet fluorescence microscopy

The cleared samples are acquired with the ultramicroscope I (LaVision BioTec-Miltenyi) assisted by the ImspectorPro software (LaVision BioTec-Miltenyi). The light sheet is created by two cylindrical lenses using laser with 3 specific wavelengths: 561nm, 640nm and 785nm (Coherent Sapphire Laser, LaVision BioTec-Miltenyi). For all the acquisitions we used a binocular stereomicroscope (MXV10, Olympus) with a 2× objective (MVPLAPO, Olympus) adding a protective dipping cap with corrective lens. For detection we used a Zyla SCMOS camera (2,048 × 2,048 pixel) from Andor Oxford Instrument. The images were in 16-bit. We used for zoom magnification 3.2X (pixel sizes x,y=1.02um). Samples were dived in a quartz tank (LaVision BioTec-Miltenyi) filled with DBE to maintain the clearing. The z step size between each image was 1μm with 150ms time exposure.

### Oocyte RNA extraction

Total RNA extraction was performed as in Ref.17 from fully grown oocytes with *zona pellucida*. For the RNA-Seq analysis, three biological replicates were performed each containing 10 oocytes from two *Myo10^flox/flox^; Cre*^+^ mice (*Myo10^-/-^* full oocytes) and two *Myo10^wt/flox^; Cre^+^ (Myo10^+/-^* oocytes). For the RT-qPCR analysis, two biological replicates were made containing 30 and 24 oocytes from two *Myo10^flox/flox^; Cre^-^* mice (*Myo10^-/-^* full oocytes) and two *Myo10^wt/flox^; Cre^+^ (Myo10^+/-^* oocytes). Total RNA extraction was performed using the RNAqueous-Micro Total RNA Isolation Kit (Thermo Fisher Scientific Ref. AM1931). Briefly, oocytes were washed three times in 1X PBS, transferred to lysis buffer (supplemented with 1 μL β-mercaptoethanol for the RNA-Seq analysis) and freezed in liquid nitrogen where they were stored at −80°C for 1 h. After thawing, RNA extraction was performed as described by the manufacturer. RNAs were eluted twice in 10 μL elution buffer and treated with DNaseI.

### RNA-Seq and data analysis

#### cDNA libraries and RNA-Seq

Library preparation and Illumina sequencing were performed at the Ecole normale superieure genomic core facility (Paris, France). 5 ng of total RNA was amplified and converted to cDNA using the SMART-Seq v4 Ultra Low Input RNA kit (Clontech Ref. 634889). Afterwards, an average of 500 pg of amplified cDNA was used to prepare the libraries following Nextera XT DNA kit (Illumina Ref. FC-131-1096). Libraries were multiplexed by 6 on a high-output flow cells. A 75 bp read sequencing was performed on a NextSeq 500 device (Illumina). A mean of 26 ± 4 million passing Illumina quality filter reads was obtained for each of the 6 samples.

#### Bioinformatics analysis

The analysis was performed using the Galaxy server of ARTbio bioinformatics platform. Sequencing quality control was carried out with FastQC (Babraham Bioinformatics) (Galaxy Version v0.72+galaxy1) and MultiQC tools (v1.6). Alignment of reads to the genome of *Mus musculus* mm10 reference genome was done with Bowtie 2 (v2.3.4.2)^52^ with fast local option for soft clipping of the Nextera transposase adapter sequence. We used mm10 GTF file (Ensembl GTF Mus_musculus.GRCm38.94.chr) as annotation files. Counting of reads per gene and differential gene expression analysis were performed with featureCounts (v1.6.3+galaxy2) and DESeq2 (v2.11.40.4), respectively. Transcripts were determined as differentially expressed based on a P adj (adjusted P value with false discovery rate correction) of less than 0.05. Of note, 222 deregulated transcripts were annotated as predicted (32.4% of the deregulated transcripts, Table.1) and were predominantly pseudogenes (84.2% of deregulated predicted transcripts).

#### Gene Ontology (GO) enrichment analysis

The Analysis was performed with the PANTHER Classification System using a Fisher’s exact test with a P value with false discovery rate correction (FDR) threshold of 0.05. For GO biological process complete (Table 1), the deregulated transcripts list from the RNA-Seq analysis was used as the input list and the *Mus musculus* genome as the reference list.

### RT-qPCR

Reverse transcription of the extracted RNAs was performed using iScript Reverse Transcription Supermix (Bio-Rad Ref. 1708840). We used SsoAdvanced Universal SYBR Green Supermix (Bio-Rad Ref. 1725270) and CFX-96 (Bio-Rad) for quantitative PCR of cDNAs using the primers shown below. Two technical replicates were made for each condition. mRNA quantity was normalized to *Gapdh* and relative levels of mRNA expression were calculated with the 2-ΔΔC. The mean and standard error of mean were normalized to *Myo10*^+/-^.

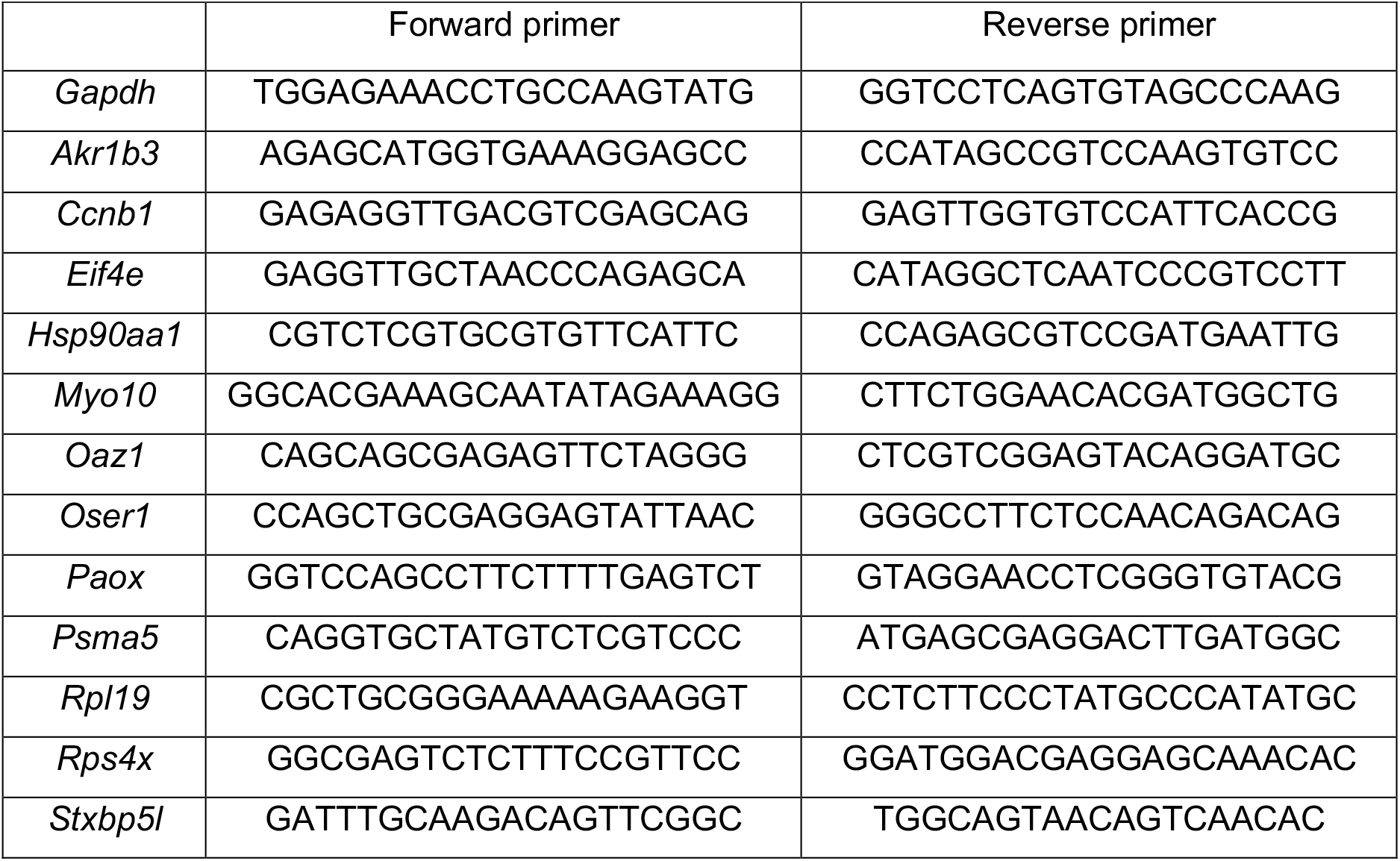

### Machine learning approach for oocyte morphological characterization

We used our machine learning pipeline presented in Ref.22 to automatically measure and compare the morphology of *Myo10*^-/-^ and control oocytes from brightfield images. Oocyte and *zona pellucida* contours were automatically determined with Oocytor neural networks. This segmentation allowed us to extract the value of 89 features for each oocyte-*zona pellucida* complex. These features characterize the size, shape, global intensity repartition, local texture and spatial organization of the oocyte, perivitelline space and *zona pellucida*. The complete list and description of the defined features are given in our github repository (https://github.com/gletort/Oocytor). As the features are not independent between themselves, we automatically selected a subset of uncorrelated features (with a threshold of 0.75) as described in Ref.22. We next trained a Random Forest classifier^53^ with these features to distinguish between fully grown *Myo10*^-/-^ and control oocytes cleared of follicular cells. Importantly for our study, the RandomForest algorithm scored each feature according to its importance for classification, based on the Gini index. We checked that the algorithm was able to recognize the oocyte type with a 10-fold cross validation on our dataset and obtained a classification accuracy of 82% (with a precision of 80.9% and a recall of 83%). The most important features for classification concerned the composition of the *zona pellucida* with the presence of vertical linear structures in it (TZP-like structure), the texture of the cytoplasm, the heterogeneity in the thickness of the perivitelline space and the circularity of both the oocyte and the internal limit of the *zona pellucida.* Manual inspection of these features suggested that the difference in the texture of the cytoplasm was mainly due to differences in nucleolus focus rather than actual morphological differences, and this feature was not statistically different between *Myo10*^-/-^ full and control oocytes during meiotic maturation. We therefore decided not to explore this characteristic further.

#### Description of the features showed in this study

##### Vertical linear structure in the zona pellucida

To quantify the presence of vertical linear structures resembling TZPs, the area in the segmented *zona pellucida* was first flattened with the « Flatten » option in Fiji (NIH) to obtain a horizontal rectangle. Linear structures were then detected by measuring and thresholding the « tubeness » of each pixel in the image (with « Tubeness » plugin in Fiji which is based on the eigenvalues of the Hessian matrix of the image). Finally, only fairly vertical structures were kept by applying Sobel filters on the thresholded images.

##### Perivitelline space thickness

Thickness was calculated as the distance between the oocyte shape and the internal limit of the *zona pellucida* at each angle around the oocyte center. The mean, minimum, maximum, standard deviation (data not shown) and coefficient of variation were then calculated from the distribution of distances.

##### Oocyte aspect ratio

Ratio between the maximum Feret diameter (longest distance) of the oocyte shape and the minimum Feret diameter (distance in the perpendicular direction to the maximum Feret diameter). A circle has an aspect ratio of 1, while an elongated shape has a higher aspect ratio.

### Image analysis

For all signal intensity measurements, the image background was subtracted using Metamorph software.

#### Signal intensity of SiR-actin labeled TZPs

Measurements were performed on growing and fully grown oocytes, cleared of follicular cells and labeled with SiR-actin. For each oocyte, the 5th image among 8 Z acquisitions spaced 4 μm apart was selected for measurement, which corresponded to the oocyte equatorial plane. After background subtraction, we drew an ellipsoidal ROI of 3 pixels (0.34 μm) width along the pericenter of the *zona pellucida* using Fiji. The mean gray value of SiR-actin signal in the ROI was measured as a readout of TZP density. For each oocyte, the mean gray value was normalized to that of the control oocyte with the highest mean gray value. The diameter of each oocyte was measured in parallel.

#### Signal intensity of EU labeled RNAs

Measurements were performed using Metamorph similarly to Ref.23 on growing and fully grown oocytes. After background subtraction and Sum Z projection of the DAPI and EU signal images (images were spaced 1 μm apart and selected for Z projection based on DAPI signal), the integrated intensity of the DAPI and EU signals was measured in a ROI of 77,577 pixels (999 μm^2^). The same ROI was used for all measurements and was drawn to encompass the DAPI signal of each oocyte. The integrated intensity of the EU signal was then normalized to that of the DAPI signal. For each oocyte, the relative EU intensity value was normalized to that of the control oocyte with the highest relative EU intensity value.

#### Signal intensity of phalloidin labeled cytoplasmic F-actin

Measurements were done on oocytes fixed 6 hours after NEBD using Metamorph as in Ref.17. After background subtraction, the integrated intensity of the phalloidin signal was measured in a ROI containing the cytoplasm of the oocyte equatorial plane. For each oocyte, we used the same ROI of 216,290 pixels (2,786 μm^2^) and the integrated intensity of the phalloidin signal was normalized to that of the control oocyte with the highest value of phalloidin signal intensity.

#### Ovarian follicular density

Cleared ovaries were imaged by light-sheet fluorescence microscopy and ovarian slices, one every 50 slices, were selected for measurement (slices were spaced 1 μm apart). For each image, the area of the ovarian slice was measured by thresholding the TO-PRO-3 signal using Fiji. Briefly, images were preprocessed with a gamma and a median filter and thresholded with the Otsu dark method. When image thresholding led to more than one ovarian area due to discontinuous signal, only the largest thresholded area was selected. For each image, we counted the number of follicles in the thresholded ovarian slice. Follicles were determined based on a defined outline, with an apparent oocyte surrounded by visible follicular cell layers (this includes preantral and antral follicles). The number of follicles in the thresholded ovarian slice was then normalized to the slice area. The follicular density of the ovary corresponds to the mean of the follicular density of the slices analyzed.

#### Timing of polar body extrusion

The timing of the first polar body extrusion was determined in oocytes monitored by live spinning disk microscopy. Only movies with control oocytes extruding a polar body before 12 hours after NEBD were selected.

#### Spindle bipolarization and positioning

Spindle morphogenesis and positioning were assessed in SiR-tubulin labeled oocytes monitored during meiotic maturation by live spinning disk microscopy. The timing of spindle bipolarization was assessed by scoring the percentage of oocytes with a bipolarized spindle every hour from 1 to 6 h after NEBD. Spindle interpolar length and central width were measured 7 h after NEBD using Metamorph. Only spindles parallel to the acquisition plane were measured. Spindle localization was assessed by scoring the percentage of oocytes with a central or off-centered spindle 30 min before anaphase for oocytes that extruded a polar body or up to 12 h after NEBD for those that did not.

#### Chromosome alignment

Metaphase plate height and width were measured as in Ref.17 using Metamorph in H2B-GFP expressing oocytes monitored during meiotic maturation by live spinning disk microscopy. We measured the height and width of a ROI rectangle enclosing chromosomes 30 min before anaphase for oocytes that extruded a polar body or 12 h after NEBD for those that did not. Only metaphase plates oriented parallel to the acquisition plane were measured.

### Statistical analysis

We used GraphPad Prism 7.0a software for statistical analysis of RT-qPCR, oocyte polar body extrusion rate and image data. The Gaussian distribution of values was tested using a D’Agostino & Pearson normality test. Comparison of means was performed by a two-tailed unpaired t test (with Welch’s correction for unequal variances) or by a two-tailed Mann Whitney test (for a non-Gaussian distribution of values). We used two-sided Fisher’s exact tests and a Chi-square test for contingency table analysis. All tests were performed with a confidence interval of 95%. n.s (not significant) P value ≥ 0.05, * P value < 0.05, ** P value < 0.01, *** P value < 0.001 and **** P value < 0.0001. For each graph, the error bars indicate the standard deviation or standard error of mean.

## Ethical statement

All experimental procedures used for the project have been approved by the ministry of agriculture to be conducted in our CIRB animal facility (authorization N°75-1170). The use of all the genetically modified organisms described in this project has been granted by the DGRI (Direction Générale de la Recherche et de l’Innovation: Agrément OGM; DUO-5291).

## SUPPLEMENTARY FIGURE LEGENDS

**Figure S1:**
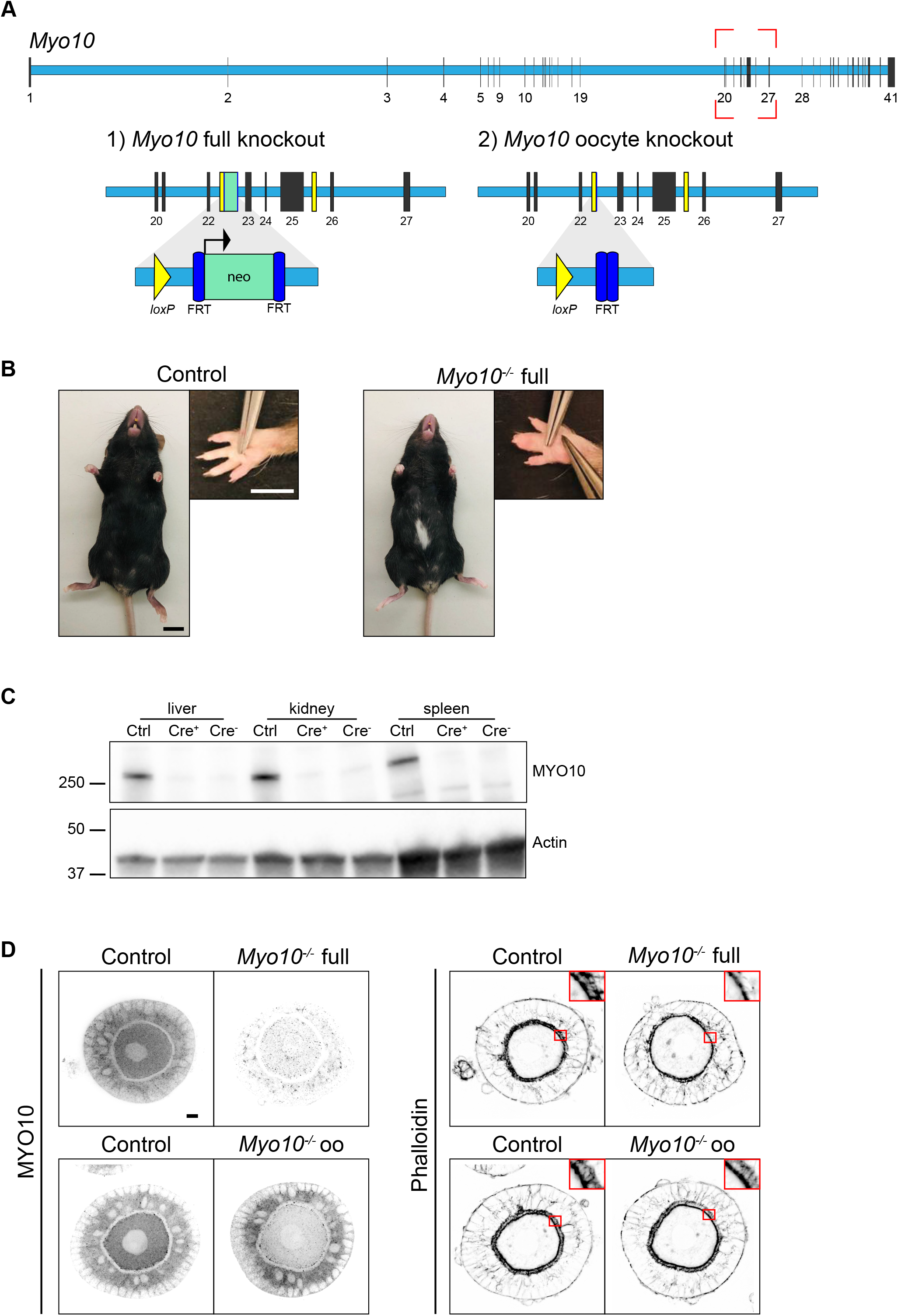
*Myo10* full knockout, generation and validation. **A)** Scheme of the *Myo10* gene with introns in light blue rectangles and exons in black vertical lines according to Ensembl database. The red dotted rectangle indicates the region targeted to invalidate MYO10. 1) For *Myo10* full knockout: the neomycin selection cassette (neo, green rectangle) inserted between exons 22 and 23 was retained, allowing its promoter to interfere with MYO10 expression in all cell types. 2) For *Myo10* conditional knockout in oocytes: the neo cassette was removed following Flp-mediated excision at the FRT sites (dark blue rectangles). ZP3 Cre-mediated excision at the *loxP* sites (yellow rectangles and triangles) flanking exons 23-25 results in a premature stop codon and absence of MYO10 in oocytes. **B)** Control (left panel) and *Myo10*^-/-^ full (right panel) mice of the *Myo10* full knockout strain. Scale bar 10 mm. For each panel, the image on the right shows the mouse front paw (Scale bar 5 mm). Some phenotypes characteristic of previously described *Myo10* full knockout were observed in our *Myo10*^-/-^ full mice, such as a white belly spot and webbed digits. **C)** Immunoblotting performed on liver, kidney and spleen extracts from mice from the *Myo10* full knockout strain. Control (Ctrl) extracts are from a *Myo10^wt/flox^; Cre*^+^ genotyped mouse, Cre^+^ extracts are from a *Myo10^flox/flox^; Cre*^+^ genotyped mouse and Cre^-^ extracts are from a *Myo10^flox/flox^; Cre^-^* genotyped mouse. The membrane was stained for MYO10 (upper image) and for Actin (lower image) as a loading control. Indicated ladders: 37, 50 and 250 kDa. **D)** Preantral follicles from the *Myo10* full knockout strain (upper images) and the *Myo10* oocyte knockout strain (lower images). Follicles were stained for MYO10 (left panel) and with phalloidin (right panel, top corners show the magnification of the red rectangles focusing on the *zona pellucida*). For each panel, control follicles are on the left and *Myo10*^-/-^ follicles on the right. For MYO10 staining, contrast adjustment is similar between follicles of the same strain. Scale bar 10 μm.

**Figure S2:**
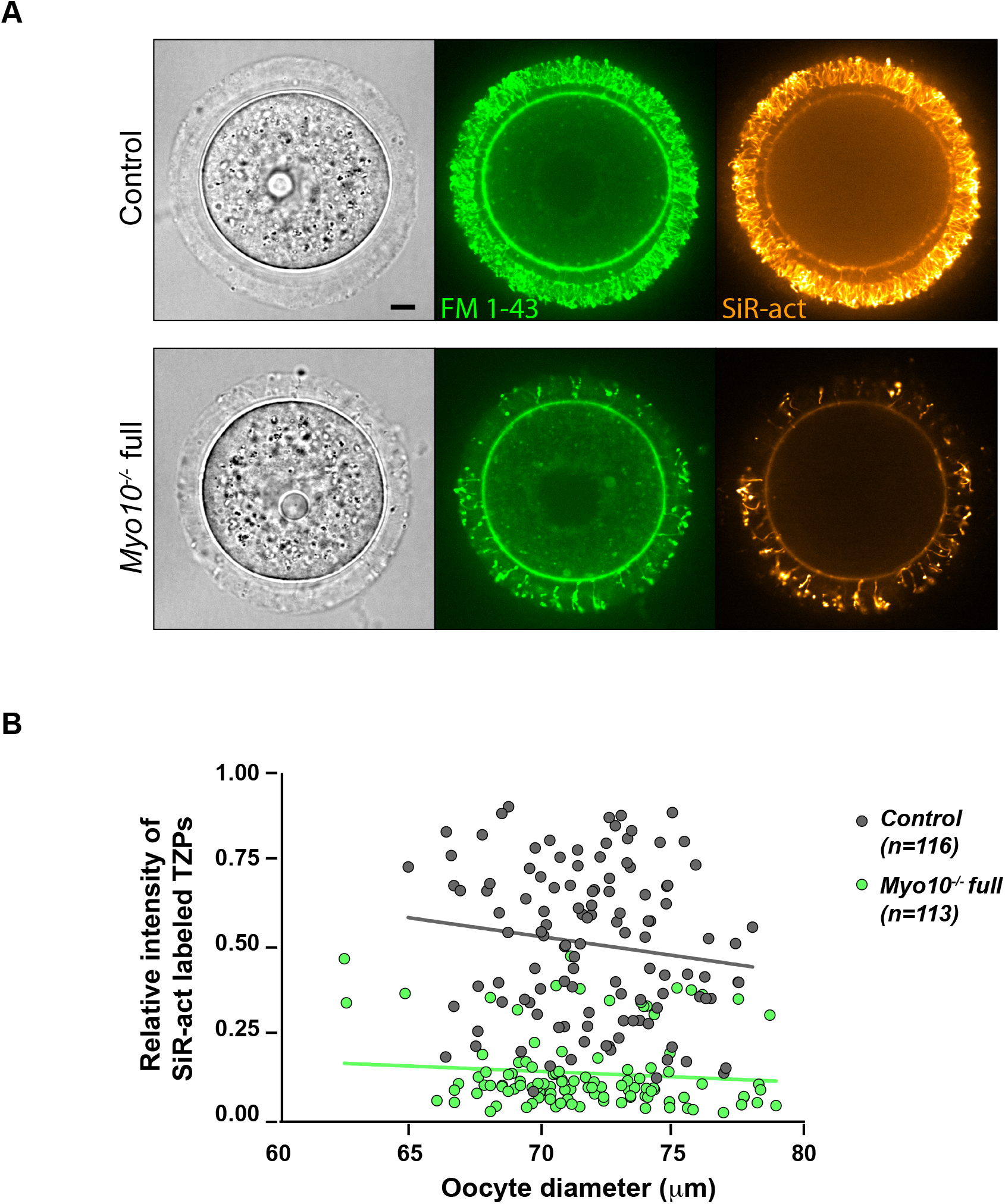
Global deletion of *Myo10* decreases the density of all transzonal projections. **A)** Images of fully grown control (upper panel) and *Myo10*^-/-^ full (lower panel) oocytes stained with FM 1-43 (green, middle images) to label membranes and SiR-actin (SiR-act, orange, right images) to label F-actin. The left images show oocyte brightfield images. Scale bar 10 μm. **B)** Scatter plot of the intensity of all SiR-act labeled TZPs (y axis) versus oocyte diameter in μm (x axis) of oocytes recovered from growing and fully grown follicles. Control and *Myo10*^-/-^ full oocytes are in dark gray and green, respectively. (n) is the number of oocytes analyzed. Data are from eight independent experiments. Lines are a simple linear regression for easier visualization.

**Figure S3:**
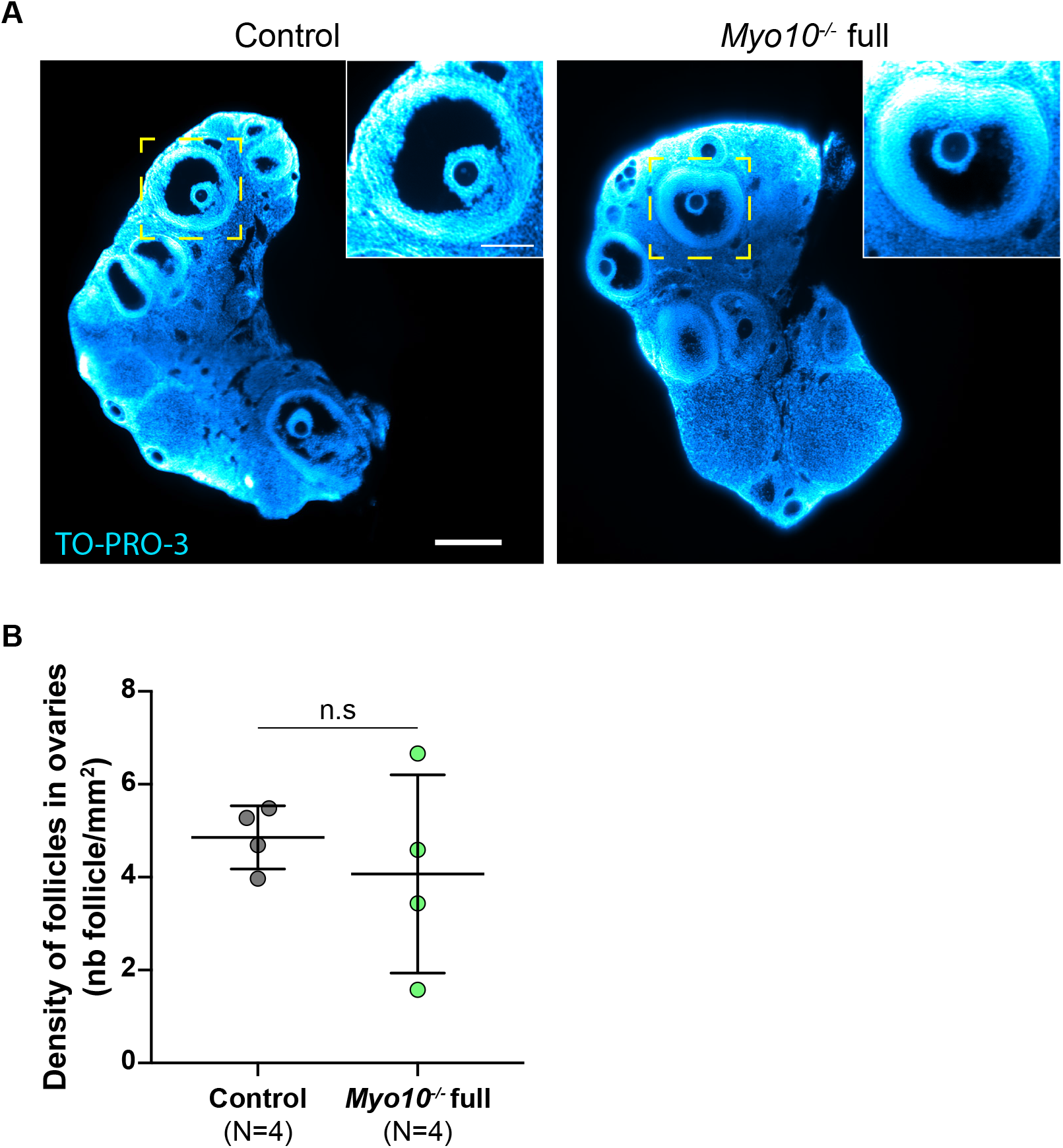
Follicular density is normal in *Myo10*^-/-^ full ovaries. **A)** Light-sheet fluorescence microscopy images of cleared ovaries stained with TO-PRO-3 (blue) to label nucleic acid. The control ovary is on the left and the *Myo10*^-/-^ full ovary on the right. Scale bar 0.25 mm. The images inserted at the top right show antral follicles from the magnification of the yellow dotted rectangles. Scale bar 0.1 mm. **B)** Scatter plot of the density of follicles in ovaries (number of follicles per mm^2^). Control ovaries are in dark gray and *Myo10*^-/-^ full ovaries in green. For each condition, data are from two different mice. (N) is the number of ovaries analyzed. Data are mean±s.d. with individual data points plotted. Statistical significance of differences was assessed by a two-tailed Mann Whitney test, P value = 0.4857. n.s not significant.

**Figure S4:**
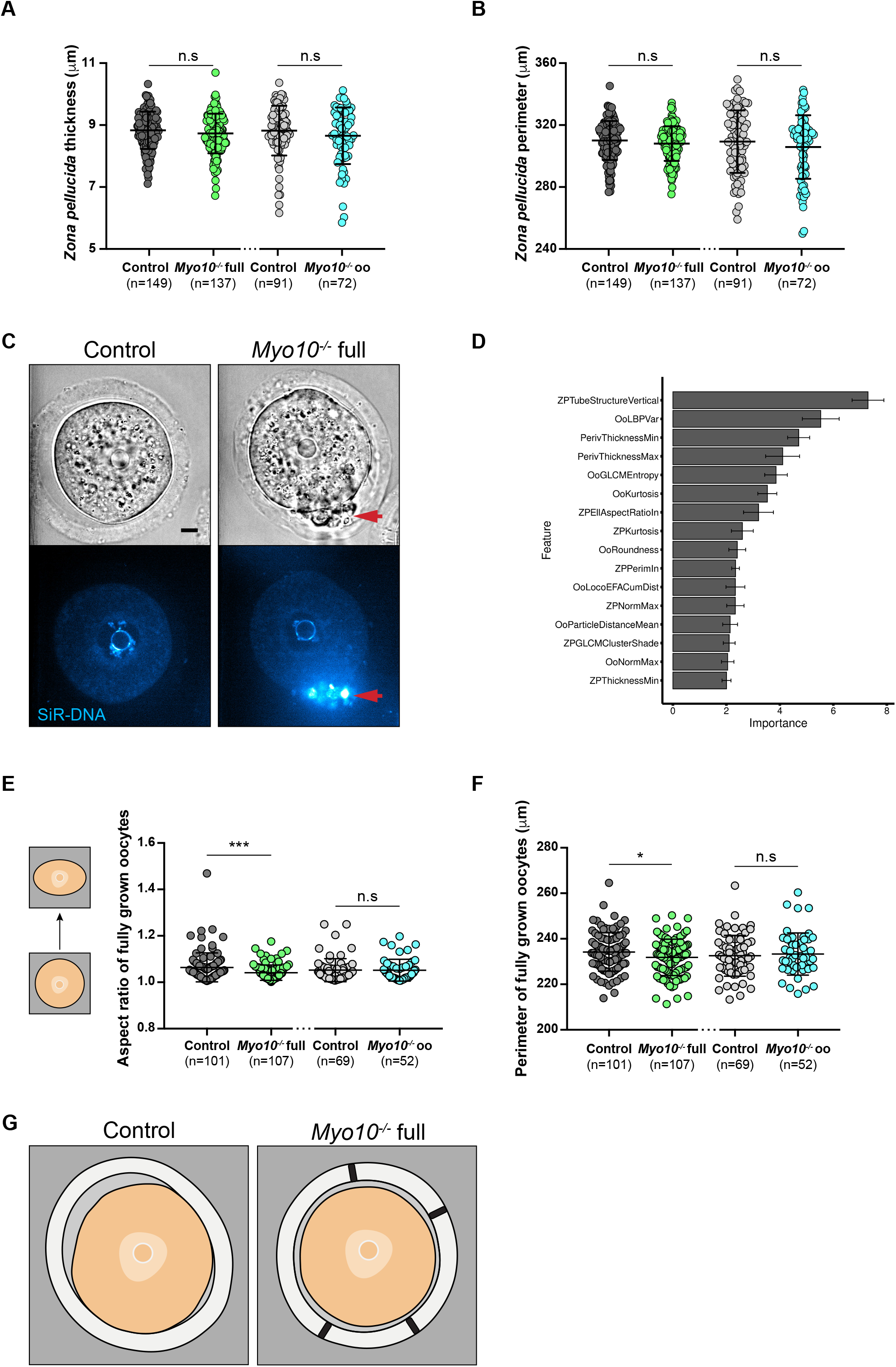
TZP-deprived oocytes have morphological differences. **A)** Scatter plot of the *zona pellucida* thickness of control and *Myo10*^-/-^ full oocytes (in dark gray and green, respectively) and control and *Myo10*^-/-^ oo oocytes (in light gray and blue, respectively). (n) is the number of oocytes analyzed. Data are mean±s.d. with individual data points plotted. Data are from four to thirteen independent experiments. Statistical significance of differences was assessed by a two-tailed unpaired t test, P value = 0.1560 (*Myo10*^-/-^ full oocytes compared to control oocytes) and by a two-tailed Mann Whitney test, P value = 0.3229 (*Myo10*^-/-^ oo oocytes compared to control oocytes). n.s not significant. **B)** Scatter plot of the *zona pellucida* perimeter of control and *Myo10*^-/-^ full oocytes (in dark gray and green, respectively) and control and *Myo10*^-/-^ oo oocytes (in light gray and blue, respectively). (n) is the number of oocytes analyzed. Data are mean±s.d. with individual data points plotted. Data are from four to thirteen independent experiments. Statistical significance of differences was assessed by two-tailed Mann Whitney tests, P value = 0.1466 (*Myo10*^-/-^ full oocytes compared to control oocytes) and P value = 0.2804 (*Myo10*^-/-^ oo oocytes compared to control oocytes). n.s not significant. **C)** Fully grown control (left panel) and *Myo10*^-/-^ full (right panel) oocytes stained with SiR-DNA (blue). For each panel, the brightfield image of the oocyte is at the top and the corresponding SiR-DNA labeling at the bottom. Some follicular-like cells are ectopically located in the perivitelline space of the *Myo10*^-/-^ full oocyte (red arrows). Scale bar 10 μm. **D)** Bar graph of the 16 most important morphological features describing oocytes used by our machine learning algorithm to discriminate control from *Myo10*^-/-^ full oocytes. Features are ranked according to their importance for oocyte classification. **E)** Scatter plot of the aspect ratio (circularity) of oocytes, as represented by the schemes on the left. Control and *Myo10*^-/-^ full oocytes are in dark gray and green, respectively. Control and *Myo10*^-/-^ oo oocytes are in light gray and blue, respectively. (n) is the number of oocytes analyzed. Data are mean±s.d. with individual data points plotted. Data are from four to thirteen independent experiments. Statistical significance of differences was assessed by two-tailed Mann Whitney tests, P value = 0.0004 (*Myo10*^-/-^ full oocytes compared to control oocytes) and P value = 0.8235 (*Myo10*^-/-^ oo oocytes compared to control oocytes). n.s not significant. **F)** Scatter plot of the perimeter of control and *Myo10*^-/-^ full oocytes (in dark gray and green, respectively) and control and *Myo10*^-/-^ oo oocytes (in light gray and blue, respectively). (n) is the number of oocytes analyzed. Data are mean±s.d. with individual data points plotted. Data are from four to thirteen independent experiments. Statistical significance of differences between *Myo10*^-/-^ full oocytes and control oocytes was assessed by a two-tailed unpaired t test, P value = 0.0409. Statistical significance of differences between *Myo10*^-/-^ oo oocytes and control oocytes was assessed by a two-tailed Mann Whitney test, P value = 0.7405. n.s not significant. **G)** Schemes summarizing the most important morphological features used by our machine learning algorithm to discriminate a control (left image) from a *Myo10*^-/-^ full oocyte (right image). Features include *zona pellucida* texture (vertical TZP-like structures passing through it, Fig.2 F, G), heterogeneity in perivitelline space thickness (Fig.2 D, E) and oocyte circularity (C).

**Figure S5:**
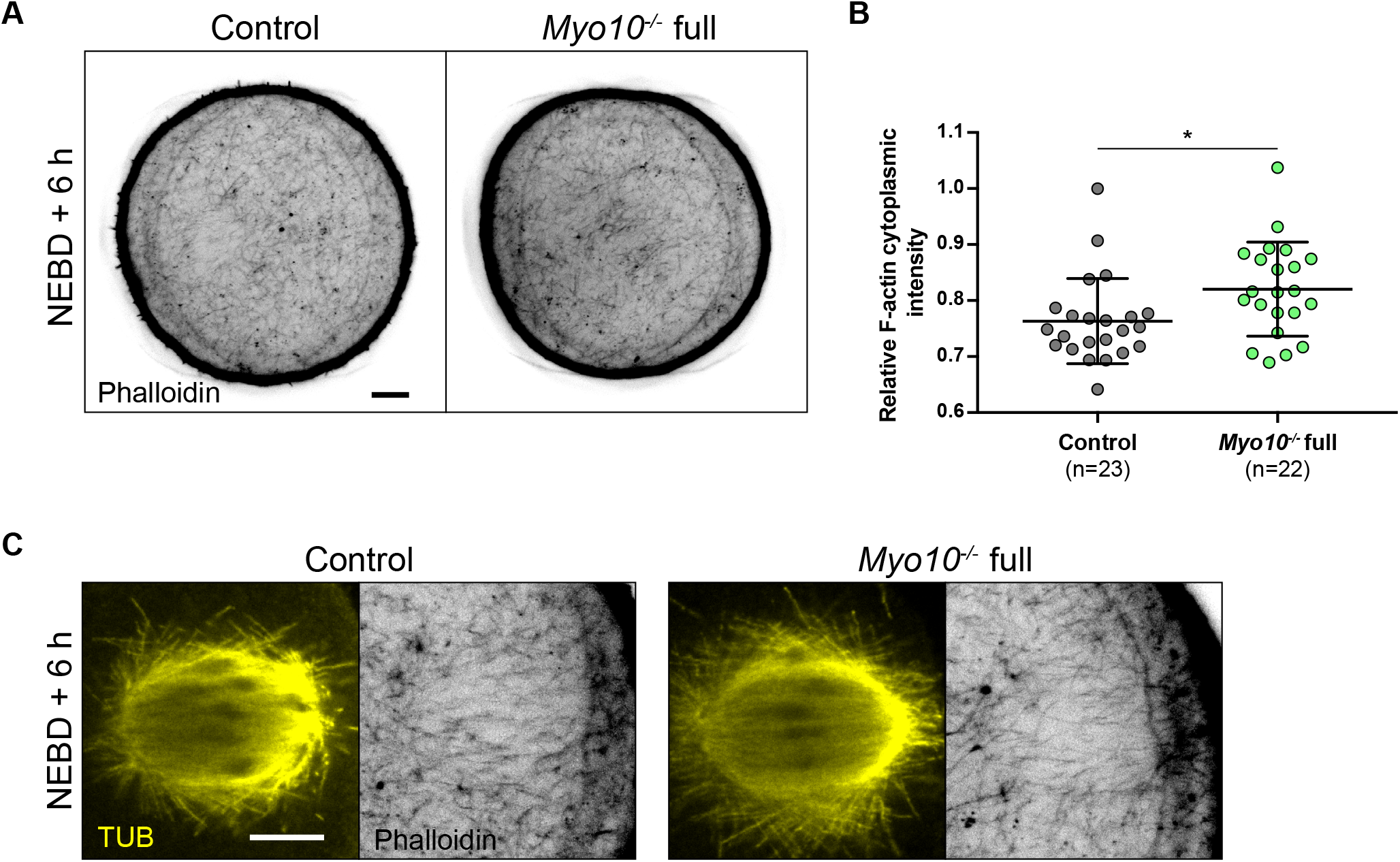
The cytoplasmic F-actin network is normally organized in TZP-deprived oocytes. **A)** Equatorial planes of control (left) and *Myo10*^-/-^ full (right) oocytes stained with phalloidin 6h after NEBD. Scale bar 10 μm. **B)** Scatter plot of cytoplasmic F-actin intensity 6h after NEBD in control (dark gray) and *Myo10*^-/-^ full (green) oocytes. (n) is the number of oocytes analyzed. Data are mean±s.d. with individual data points plotted. Data are from three independent experiments. Statistical significance of differences was assessed by a two-tailed Mann Whitney test, P value = 0.0111. **C)** Images of spindles and F-actin cages from control (left panel) and *Myo10*^-/-^ full (right panel) oocytes 6h after NEBD. Oocytes are stained for Tubulin (TUB, yellow, left panel images) and stained with phalloidin (black, right panel images). Scale bar 10 μm.

---

<u>**Table 1: Differential gene expression between fully grown *Mvo10*^+/-^ and *Myo10*^-/-^ oocytes performed by RNA-Seq**</u>

**Sheet 1)** Raw data from the RNA-Seq analysis shown in Fig.3 A.

**Sheet 2)** The 685 deregulated transcripts selected with a P adj < 0.05. The transcripts highlighted in green are those selected to be validated by the RT-qPCR analysis presented in Fig.3 C.

**Sheet 3)** The exhaustive list of significantly over-represented GO biological process terms in the list of deregulated transcripts. Significance was set at FDR P < 0.05.

**Movie S1: Decreased TZP density in *Myo10*^-/-^ full oocytes highlighted by OMX super-resolution microscopy**

OMX microscopy movies showing TZPs of fully grown oocytes freed of follicular cells and stained with phalloidin. The control oocyte is on the left and the *Myo10*^-/-^ full oocyte on the right. Movies are 3D projection images reconstructed from the 125 nm spaced Z acquisition from the outer layer of the *zona pellucida* (ZP) to the oocyte cortex. Movies start with the top view of the ZP outer layer, rotate to a ZP cross-sectional view and come back to the ZP outer layer. Scale bar 5 μm.

**Movie S2: Intra-ovarian organization of *Myo1Q*^-/-^ full ovaries**

Light-sheet fluorescence microscopy movies of cleared ovaries stained with TO-PRO-3 to label nucleic acid. Movies are cross-sectional images of the ovaries spaced 1 μm apart. **A)** Movies of four ovaries from two adult control mice. **B)** Movies of four ovaries from two adult *Myo10*^-/-^ full mice. Scale bars 200 μm.

**Movie S3: Some TZP-deprived oocytes arrest in meiosis I**

Spinning disk movies showing brightfield images of a control oocyte (top movie), a *Myo10*^-/-^ full oocyte that extruded a polar body (middle movie) and a *Myo10*^-/-^ full oocyte that did not extrude a polar body (bottom movie). The oocytes are those shown in Fig.4 A. Movies start 2 h 30 after NEBD and end 18 h after NEBD. Images are acquired every 30 min. Scale bar 10 μm.

**Movie S4: TZP-deprived oocytes correctly form and position meiosis I spindles**

Spinning disk movies of oocytes stained with SiR-tubulin (SiR-tub) to label microtubules (yellow) from 30 min to 14 h 30 after NEBD. The top movies show a control oocyte, the middle ones a *Myo10*^-/-^ full oocyte that extruded a polar body and the bottom ones a *Myo10*^-/-^ full oocyte that did not extrude a polar body. For each oocyte, the left movie shows Max intensity Z Projection of SiR-tub labeling and the right one the same SiR-tub labeling merged with the corresponding brightfield images. The oocytes are those shown in Fig.5 A. Images are acquired every 30 min. Scale bar 10 μm.

**Movie S5: TZP-deprived oocytes properly align meiosis I chromosomes**

Spinning disk movies of oocytes injected with H2B-GFP to label chromosomes from 2 h 30 to 18 h after NEBD. The top movies show a control oocyte, the middle ones a *Myo10*^-/-^ full oocyte that extruded a polar body, and the bottom ones a *Myo10*^-/-^ full oocyte that did not extrude a polar body. For each oocyte type, the left movie shows Max Intensity Z Projection of H2B labeling and the right one the same H2B labeling merged with the corresponding brightfield image. The oocytes are those shown in Fig.5 F. Images are acquired every 30 min. Scale bar 10 μm.

